# N^6^-Methyladenine DNA Modification in Human Genome

**DOI:** 10.1101/176958

**Authors:** Chuan-Le Xiao, Song Zhu, Minghui He, De Chen, Qian Zhang, Ying Chen, Guoliang Yu, Jinbao Liu, Shang-Qian Xie, Feng Luo, Zhe Liang, De-Peng Wang, Xiao-Chen Bo, Xiao-Feng Gu, Kai Wang, Guang-Rong Yan

## Abstract

DNA N6-methyladenine (6mA) modification is the most prevalent DNA modification in prokaryotes, but whether it exists in human cells and whether it plays a role in human diseases remain enigmatic. Here, we showed that 6mA is extensively present in human genome, and we cataloged 881,240 6mA sites accounting for ∼0.051% of the total adenines. [G/C]AGG[C/T] was the most significantly associated motif with 6mA modification. 6mA sites were enriched in the coding regions and mark actively transcribed genes in human cells. We further found that DNA N6-methyladenine and N6-demethyladenine modification in human genome were mediated by methyltransferase N6AMT1 and demethylase ALKBH1, respectively. The abundance of 6mA was significantly lower in cancers, accompaning with decreased N6AMT1 and increased ALKBH1 levels, and down-regulation of 6mA modification levels promoted tumorigenesis. Collectively, our results demonstrate that DNA 6mA modification is extensively present in human cells and the decrease of genomic DNA 6mA promotes human tumorigenesis.

## Introduction

DNA methylation is a crucial component of epigenetic regulation and plays an important role in regulating genomic imprinting, X-chromosome inactivation, transposon suppression, gene expression, epigenetic memory maintenance, embryonic development and human tumorigenesis (Bergman and Cedar, 2013; Smith and Meissner, 2013; von Meyenn et al., 2016). The most predominant DNA methylation modification in eukaryotes is 5-methylcytosine (5mC). Previous research has focused on characterizing 5mC due to its abundance and significance in eukaryotes. In contrast, N6-methyladenine DNA (6mA) modification is the most prevalent DNA modification in prokaryotes. 6mA plays crucial roles in the regulation of restriction-modification (R-M) system, DNA replication and mismatch repair, nucleoid segregation, gene expression, and is essential for viability of some bacterial strains (Luo et al., 2015; Vasu and Nagaraja, 2013; Wion and Casadesus, 2006).

DNA 6mA modification used to be considered as absent in eukaryotes including humans since it was not detectable in earlier generations of studies (Ratel et al., 2006). Recently, with the developments of deep-sequencing, 6mA was found to be present in a limited number of eukaryotes including *Chlamydomonas reinhardti* (Fu et al., 2015), *Caenorhabiditis elegans* (Greer et al., 2015), *Drosophila melanogaster* (Zhang et al., 2015), fungi (Mondo et al., 2017), mouse (Wu et al., 2016), zebrafish and pig (Liu et al., 2016b). Some studies demonstrated that 6mA participated in the regulation of gene and transposon expression including activation and suppression (Fu et al., 2015; Wu et al., 2016; Zhang et al., 2015). 6mA has also been identified as an epigenetic mark that carries heritable epigenetic information in *C. elegans* (Greer et al., 2015). Although RNA m6A modification is known to widely occur in human mRNAs and play crucial roles in RNA splicing, mRNA stability and gene expression (Wojtas et al., 2017; Xiang et al., 2017), DNA 6mA modification is considered to be absent in human genome, since it was not detectable in previous studies (Ratel et al., 2006). While many research studies have focused on 5mC modification in human, the issue of whether DNA 6mA modification occurs in general and whether it plays a role in gene regulation and disease pathogenesis remain largely unexplored.

In this study, we used a new sequencing technology, the PacBio single-molecule real-time (SMRT) sequencing, to decode and identify the presence of 6mA in human genomic DNA, especially in the mitochondria genome (Flusberg et al., 2010; Ye et al., 2017). We found that 6mA was broadly distributed across the human genome and [G/C]AGG[C/T] was the most prevalent motif at the 6mA modification sites. 6mA density was significantly enriched in exonic regions and was associated with gene transcriptional activation. Furthermore, we identified N6AMT1 and ALKBH1 as the methyltransferase and demethylase for 6mA modification in human, respectively, and we explored the potential function and clinical significance of 6mA in human tumorigenesis.

## Results

### DNA 6mA modification occurs in human genome

Using PacBio sequencing data from HuaXia1 (HX1) human genome (103х genome-wide coverage) (Shi et al., 2016), we identified 881,240 6mA modification sites, whose density (6mA/A) was approximately 0.051% of the total adenines in the human genome (See also Table S1 and S2). We found that 6mA density in human was lower than that in *fungi* (0.048%-2.8%) (Mondo et al., 2017), *Chlamydomonas* (∼0.4%) (Fu et al., 2015) and *C. elegans* (∼0.7%) (Greer et al., 2015), while similar to that in Drosophila (∼0.07% at early embryo stage) (Zhang et al., 2015).

We further applied 6mA-IP-seq to validate the presence of 6mA modification in human genomic DNA. We used a 6mA-specific antibody to enrich the 6mA-containing DNA fragments from the human blood samples, and the immunoprecipitated DNA fragments were subjected to high-throughput sequencing. Despite the relatively low sensitivity of this approach, we identified 21,129 high-confidence 6mA peaks in human blood genomic DNA (See also Table S3), confirming the presence of 6mA modifications. We further demonstrated that the 6mA-containing DNA regions identified by 6mA-IP-seq were highly overlapped with the 6mA site-occupied regions identified by SMRT sequencing (Fig. S1).

To further validate the level of 6mA in human genomic DNA by an orthogonal technology, we applied an LC-MS/MS assay on the same blood-derived DNA sample using the pure 6mA nucleoside (Berry& Associates PR 3740) as an external standard. The 6mA modification in blood sample was identified only when both its retention time and fragmentation pattern matched with 6mA standard. Our results showed that the 6mA/A ratio was ∼0.056% in the blood genome DNA sample, consistent with the ratio resolved by SMRT sequencing (∼0.051%). As SMRT sequencing approach does not discriminate between 6mA and 1mA, we further performed LC-MS/MS assays using the pure 1mA nucleoside as an external standard (Berry& Associates PR 3032). We found that 1mA was not detectable in blood genomic DNA sample used in this study (See also Fig. S2A), suggesting that SMRT sequencing was suitable for detecting 6mA in this study.

To further confirm the reliability of the identified 6mA sites in human genome from SMRT sequencing, we used 6mA-IP-qPCR technology to investigate the 6mA modification of each 10 highly, lowly and undetectably 6mA-methylated gene loci from SMRT sequencing data (See also Table S4). The results showed that the gene loci with high 6mA degree had the high enrichment of 6mA modification, the loci with low 6mA degree had low enrichment, and the loci without 6mA modification had almost no enrichment (See also Fig. S2B), suggesting that the database of 6mA sites in human genome from SMRT sequencing was reliable. Collectively, our results indicated that DNA 6mA modification was present in human genome, and we compiled a list of 6mA modification sites in human blood samples.

### 6mA density is low in chromosome X and Y, but high in mitochondria

A broad distribution of the 6mA sites across autosomal chromosomes (0.050%-0.064%) was observed in the human genome, but approximately half the density was observed in chromosomes X (0.023%) and Y (0.024%) (Fig. 1A), where it is known to have higher levels of DNA 5mC modifications. Previous study also demonstrated a strong negative correlation between the presence of 6mA epigenomic mark and 5mC epigenomic mark in *fungi* (Mondo et al., 2017). Additionally, a higher 6mA density (0.184%) was observed in mitochondria genome than that in autosomal chromosomes (Fig. 1A).

**Figure 1.**
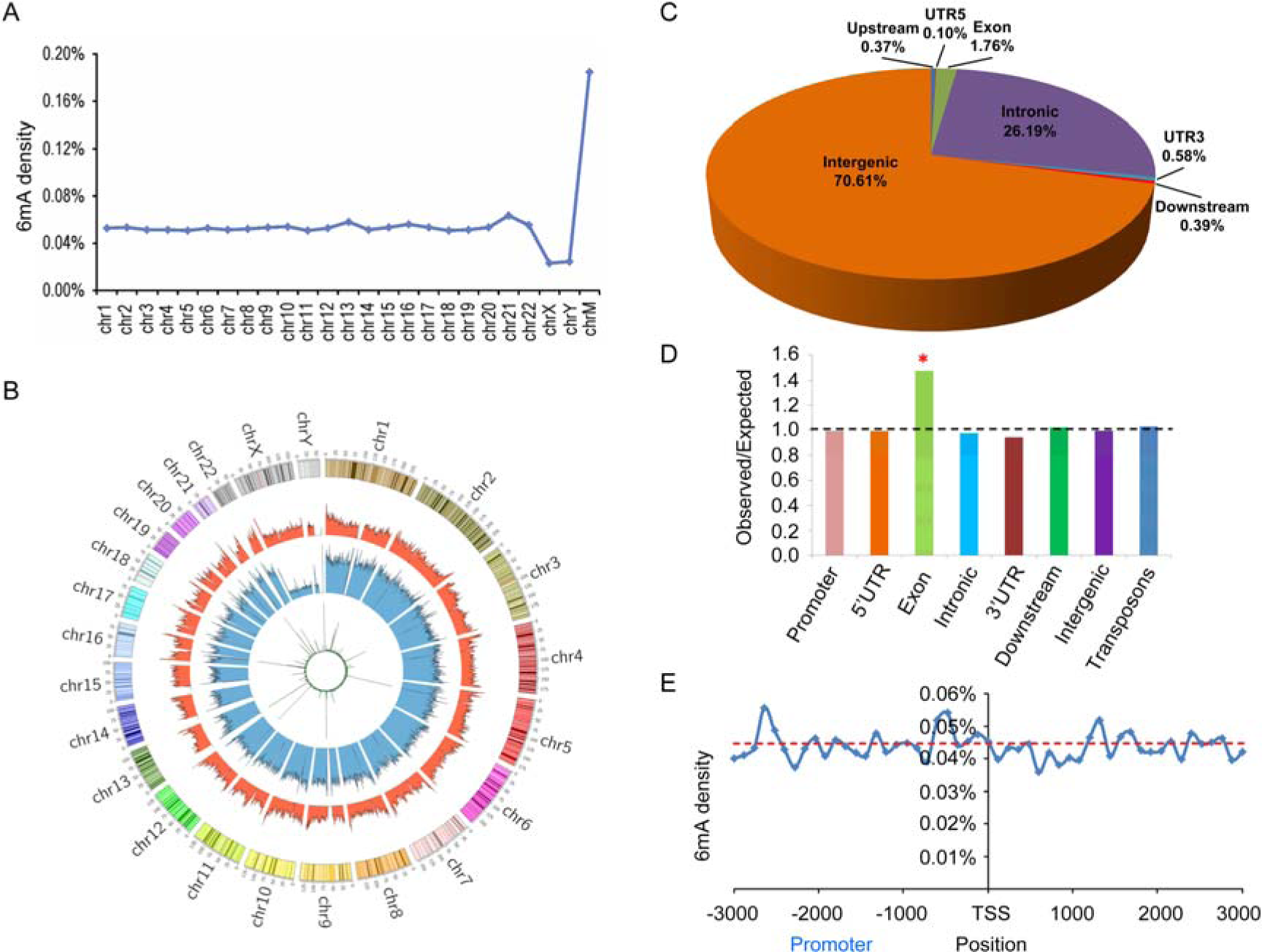
Distribution of 6mA sites in the human genome. (A) 6mA modification density across all chromosomes. (B) Circos plots of the density distribution of 6mA across all human chromosomes in the different 6mA modification level category. Green, blue, and orange circle represents lowly methylated (0-30%), moderately methylated (30%-70%), and highly methylated (70%-100%) 6mA, respectively. (C) The 6mA modification distribution in the functional elements of human genomic DNA. (D) Comparison of observed versus expected distributions of 6mA modification in each functional element. (E) Distribution of 6mA density surrounding transcriptional start sites (TSS). See also Figure S1, S2 and S3 and Table S1, S2 and S3.

We divided the 6mA modification level into low (0-30%, green circle), middle (30%-70%, blue circle), high (70%-100%, orange circle) categories and presented the data in Circos plot, in which concentric rings represent the density distribution of 6mA across all human chromosomes in the given category. We found the 6mA density was enriched in the middle 6mA modification level category (Fig. 1B), indicating that the middle 6mA level was more variable across the genome. Moreover, the 6mA density in the sexual chromosomes was not different with that in autosomal chromosomes when 6mA modification level was high. We also found a lower 6mA modification density across the centromeres, where there were many repeat elements and gaps that were not filled in the reference genome (Fig. 1B). Importantly, the chromosomal distributions of the 6mA-containing DNA fragments identified by 6mA-IP-seq were similar with those identified by SMRT sequencing despite the different sensitivity of the technologies (See also Fig. S3A and S3B).

### 6mA is significantly enriched in exon regions

Different genes may vary greatly in their 6mA density (See also Table S2). Mapping genomic distributions of 6mA modification can be an effective first step for uncovering potential functions of DNA 6mA modification (Jones, 2012; Smith and Meissner, 2013). Therefore, we examined the distributions of 6mA on the different functional regions in the human genome. We found that most of the 6mA modification sites were located in the intron and intergenic regions of genome (26.19% and 70.61%, respectively) (Fig. 1C), similar to the findings identified by 6mA-IP-seq (See also Fig. S3C), since the intron and intergenic regions accounted for most of the human genome (>95%). We found that about 23,000 genes harbored 6mA modification sites in exon and/or intron regions. Surprisingly, we found that 6mA modification density was significantly enriched in the exon-coding regions (p=0.02 Wilcoxon test, Fig. 1D), but not in the promoter, 5’UTR, intronic, 3’UTR, downstream, intergenic and transposon regions, suggesting that 6mA modification may be involved in the regulation of gene expression (Fig. 1D). Importantly, the 6mA enrichment in exon regions discovered by SMRT sequencing was also indentified by 6mA-IP-seq despite differences in sensitivity (See also Fig. S3D).

It is well known that DNA 5mC modifications in gene promoter region play an important role in epigenetic regulation of gene transcription (Bergman and Cedar, 2013). Therefore, the distribution of 6mA modification around the upstream −3kb region and downstream 3kb region of transcription start sites (TSS) and transcription terminal sites (TTS) was further examined. However, no significant enrichment of 6mA around TSS and TTS was observed in the human genome (Fig. 1E), different from what was reported on the *C. reinhardtii* genome (Fu et al., 2015).

Collectively, our results indicated that the 6mA was highly enriched in exon regions and the distribution of 6mA in the human genome can differ from those in other prokaryotes and eukaryotes. Reminiscent of the genome distribution of 5mC, 6mA distribution patterns are thus species-specific, likely reflecting different functions and mechanisms of the DNA 6mA modification.

### 6mA modification frequently occurs at sites with the AGG motif

DNA sequence motif represents short nucleotides that is widespread and is conjectured to have a biological function such as serving as transcription factor binding sites. Most of the 6mA sites are located within palindromic sequences in prokaryotes and *T. thermophila (Touzain et al., 2011)*. The 6mA sites are mainly located at ApT motif around TSS in *Chlamydomonas* (Fu et al., 2015), at AGAA and GAGG sequence motif in C. *elegans* (Greer et al., 2015). We further investigated the enriched sequence motif of the 6mA sites in human genome. We found that there was high probability (frequency higher than 50%) of guanine (G) in upstream or downstream 1bp of 6mA modification sites, which constituted a core sequence pattern (GAG) in the sequence flanking the 6mA sites (Fig. 2A). Using MEME to further analyze the flanking region to find enriched motif sequence (Bailey et al., 2009), the [G/C]AGG[C/T] motif was the most prevalent motif in 6mA sites in the human genome (Fig. 2B), similar to the motif previously discovered in the *C. elegans* genome (Greer et al., 2015). Importantly, the common sequence in 6mA-containing DNA fragments identified by 6mA-IP-seq was similar to the AGG motif identified by SMRT sequencing (Fig. 2C). Collectively, our results indicates that [G/C]AGG[C/T] is the most significantly associated motif with 6mA modification.

**Figure 2.**
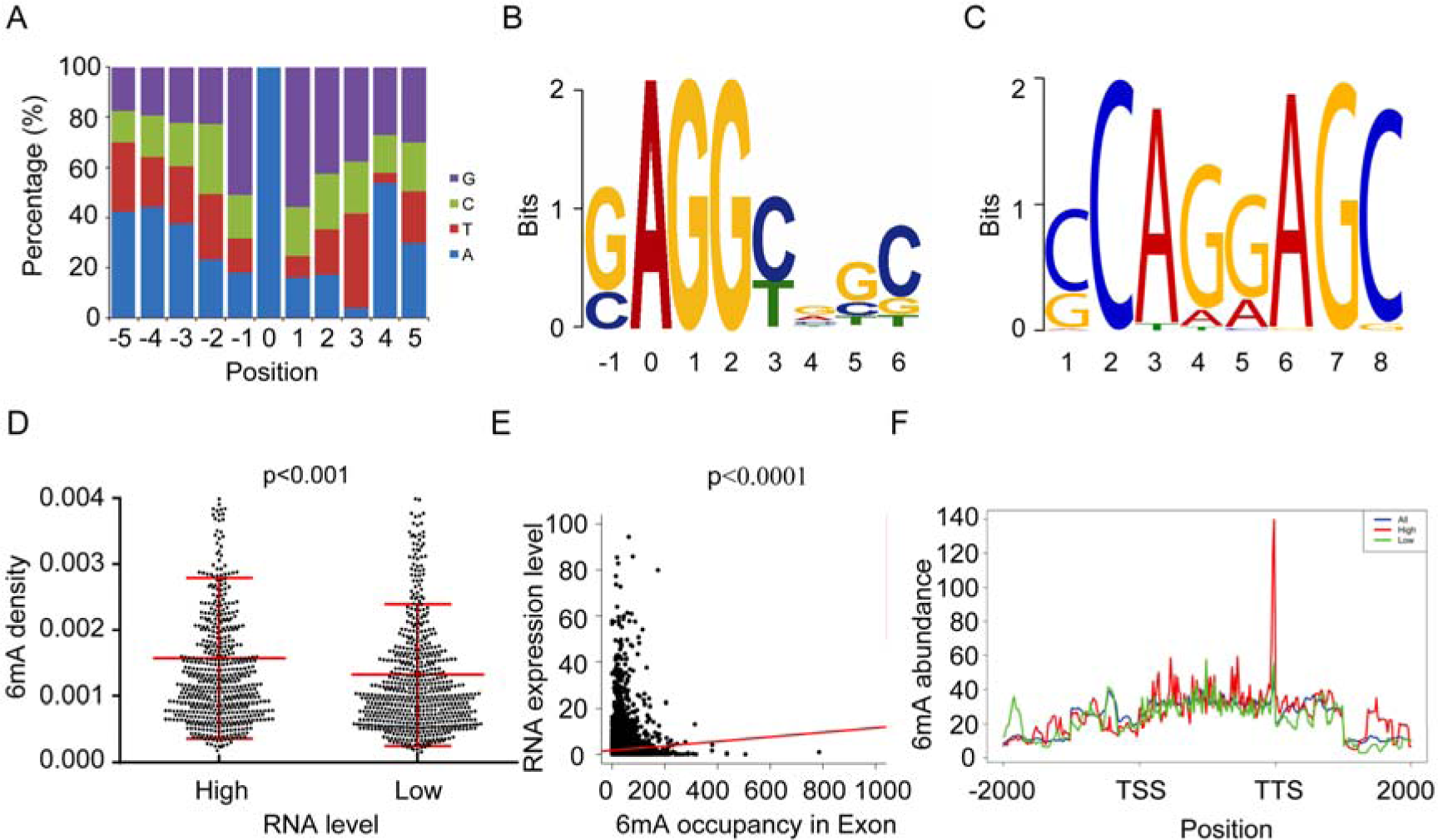
AGG is the most prevalent motif sequence of 6mA modification, and the 6mA density in exon region is significantly correlated with actively transcribed genes. (A) Base constitution percentage in the upstream and downstream flanking region of the 6mA site (“0” position). (B) The motif sequence of 6mA identify by SMRT sequencing and MEME analysis. (C) The common sequence in 6mA-containing DNA fragment peaks identified by 6mA-IP-seq. (D) Two groups of genes with top 1000 highest and top 1000 lowest expression levels (RNA-seq data (Shi et al., 2016)) are plotted with its methylation density (*p<0.001*, Student t-test). (E) The correlation between the reads of 6mA-containing DNA fragments identified by 6mA-IP-seq and RNA expression level. (F) Two groups of genes with the top 1000 highest (red) and the top 1000 lowest (green) expression levels are plotted with their 6mA abundance levels. TSS: transcription start sites; TTS: transcription terminal sites. See also Figure S4 and Table S4.

### Functional enrichment analysis on genes with higher 6mA density

To explore the possible functions of the 6mA modification, we performed GO enrichment analysis on the top 200 genes with the highest 6mA modification density in transcribed regions. Interestingly, we found that this list was significantly enriched in G-protein coupled receptor (GPCR) category (See also Fig. S4A and Table S5). Our 6mA-IP-qPCR assay confirmed experimentally that the GPCR genes had high 6mA modification ratio (See also Fig. S4B). GPCRs are well known as receptors involving in several cell signal transduction processes to alter the state of the cell. GPCRs are involved in many diseases such as cancers, and are the targets for approximately 40% of all drugs (Wacker et al., 2017). The 6mA modification of the GPCR genes may play an important role in the regulation of GPCR gene expression, but additional studies are warranted to further confirm this finding.

### Higher 6mA density in exon is positively associated with gene transcription

N6-adenine methylation modification in eukaryotic mRNA was recently shown to have profound effects on gene expression (Wojtas et al., 2017; Xiang et al., 2017). However, the potential roles of DNA 6mA modification in shaping gene expression remain largely unknown. Previous studies showed that the DNA 6mA modification in TSS activates gene expression in *Chlamydomonas* (Fu et al., 2015), while epigenetically silences gene expression in mouse embryonic stem cells (Wu et al., 2016). Therefore, we further investigated the influences of DNA 6mA modification on gene expression. RNA-seq data from HuaXia1 (HX1) human blood, which was also used for SMRT sequencing and 6mA identification, was used for this analysis (Shi et al., 2016). We selected the top 1000 highest and top 1000 lowest expressed genes as two groups and plotted their 6mA density. We found a general trend that genes with higher RNA expression tend to have higher density of 6mA around the exon regions (p<0.001 Student t-test, Fig. 2D), but not in other regions such as promoter, 5’UTR, introne, 3’UTR and intergene.

At the same time, we further investigated the correlation between the reads (abundance) of the 6mA-containing DNA fragments indentified by 6mA-IP-seq with RNA expression level. We found the positive correlation between 6mA abundance in exon regions and RNA expression level (p<0.0001) (Fig. 2E), but no correlations in other regions such as promoter, promoter, 5’UTR, introne, 3’UTR and intergene. In addition, we also selected the top 1000 highest and top 1000 lowest expressed genes as two groups and plotted their 6mA abundance. We found a general trend that genes with higher RNA expression tend to have higher 6mA abundance around the exon regions from TSS to TTS, especially around TTS sites (Fig. 2F). Taken together, our results reveal that 6mA marks the exon regions of actively transcribed genes in human cells.

### N6AMT1 is a methyltransferase for 6mA in human genome

We next focused on discovering the methyltransferase responsible for DNA 6mA modification in human. N-6 adenine-specific DNA methyltransferase 1 (putative) (N6AMT1) was originally named based on the presence of an adenine methyltransferase-characterized amino acid motif (D/N/S) PP (Y/F/W), which was a potential S-Adenosyl-l-methionine (AdoMet) dependent methyltransferase, partially similar with M.TaqI-like 6mA DNA methyltransferases of bacterial restriction-modification systems (Timinskas et al., 1995). Therefore, we speculate that N6AMT1 may be a methyltransferase for 6mA in human. To validate the hypothesis, we investigate the influences of *N6AMT1* on the 6mA level in human genome DNA *in vivo* and *in vitro*. Dot blotting and LC-MS/MS assays showed that silencing of *N6AMT1* decreased the genomics DNA 6mA modification level in human cancer cells (Fig. 3A-3C, see also Fig. S5A-S5C), while over-expression of *N6AMT1* increased the genomics DNA 6mA modification level in a dose-dependent manner (Fig. 3D-3F, see also Fig. S5D-S5F). However, silencing or over-expression of *N6AMT1* did not change RNA m6A levels (see also Fig. S6A-S6D).

**Figure 3.**
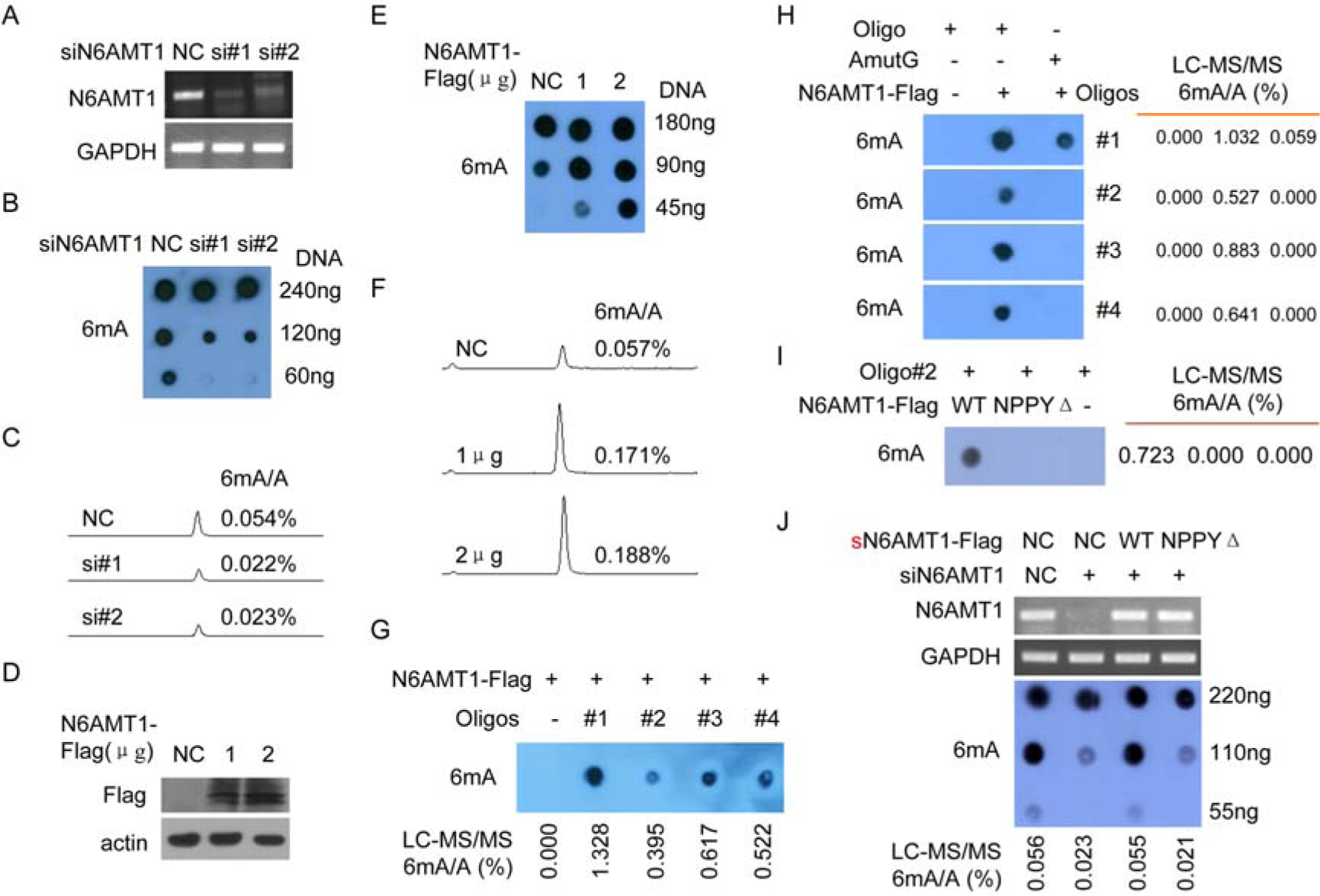
N6AMT1 is a methyltransferase for DNA 6mA methylation in human, and NPPY is the catalytic motif of N6AMT1. (A-C) MHCC-LM3 cells were transfected with anti-N6AMT1 siRNAs, the N6AMT1 levels detected by RT-PCR (A), the 6mA levels of genomic DNA were subjected to dot blotting assays using a specific anti-6mA antibody (B) and LC-MS/MS assay (C). (D-F) BEL-7402 cells were transfected with the indicated amounts of N6AMT1-Flag plasmids, the N6AMT1 levels were determined by western blotting (D), the 6mA levels of genomic DNA were subjected to dot blotting (E) and LC-MS/MS (F) assays. (G) Recombinant N6AMT1 directly methylated the DNA oligos in the *in vitro* methylation reaction. Four oligo substrates were used for 6mA-methylating by recombinant N6AMT1-Flag. (H) Recombinant N6AMT1 did not methylate the mutated DNA oligos (A mutated to G) in the *in vitro* methylation reaction. The 6mA levels were detected by dot blotting (left panel) and LC-MS/MS (right panel) assays. (I) Recombinant N6AMT1 NPPYΔ mutant did not methylate the DNA oligos compared to recombinant wild type N6AMT1 in the *in vitro* methylation reaction. (J) SK-hep1 cells were co-transfected with anti-N6AMT1 siRNAs together with the indicated synonymous mutant sN6AMT1-Flag plasmids, the N6AMT1 levels were determined by RT-PCR (upper panel), the 6mA levels were determined by dot blotting (middle panel) and LC-MS/MS (low panel) assays. See also Figure S5 and S6.

Furthermore, we performed an *in vitro* methylation assay to further determine whether N6AMT1 is a direct 6mA methyltransferase in human. The recombinant N6AMT1-Flag directly and efficiently increased the 6mA modification level from four synthetic oligonucleotide substrates (Fig. 3G), while not from the A mutation oligonucleotides of these four synthetic oligonucleotide substrates (only A was mutated to G in oligo#1 substrates which contained 23 A) (Fig. 3H).

The catalytic conserved motif NPPY was found in almost all 6mA DNA methyltransferases (MTases) in bacterial. We also found that human DNA 6mA methyltransferase N6AMT1 contains the NPPY motif (122-125aa). The recombinant N6AMT1 NPPY122-125AAAA (NPPYΔ) mutant did not directly and efficiently increase the 6mA modification level from the synthetic oligonucleotide substrates in an *in vitro* methylation assay, compared to wild-type N6AMT1 (Fig. 3I). The reduction of genomic 6mA induced by silencing of *N6AMT1* in cancer cells could be efficiently rescued by ectopic expression of wild-type but not NPPYΔ mutant *sN6AMT1* in which the *N6AMT1* sequences targeted by anti-N6AMT1 siRNA were synonymously mutated (Fig. 3J). Furthermore, we generated *N6AMT1* knockout (KO) HEK293T cell line HEK293T-N6AMT1^null^ via CRISPR-Cas9 technology (See also Fig. S7D and S7E). Dot blotting and LC-MS/MS analyses showed that genomics DNA 6mA levels were markedly decreased in N6AMT1 knockout HEK293T-N6AMT1^null^ cells (See also lane 1 and 2 in Fig. S7F). Ectopic expression of *N6AMT1* rescued the reduction of genomics DNA 6mA levels in HEK293T-N6AMT1^null^ cells, whereas the *N6AMT1* NPPYΔ mutant did not rescued the reduction of genomics DNA 6mA levels (See also Fig. S7F). Taken together, our results indicated the N6AMT1 is a methyltransferase responsible for DNA 6mA modification in human genome.

### ALKBH1 is a demethylase for DNA 6mA modification in human

The ALKBH family proteins, which contain the conserved Fe2^+^ and 2-oxoglutarate-dependent, dioxygenase domain, were promising candidates as demethylase in human (Liu et al., 2016a; Westbye et al., 2008). Among them, ALKBH2 and ALKBH3 can efficiently remove 1mA or 3mC from DNA or RNA, but not 6Ma (Aas et al., 2003; Duncan et al., 2002). We speculate ALKBH1 as a demethylase for 6mA in human given that *Homo sapiens* ALKBH1 shares high sequence homology with *Mus musculus* ALKBH1, which is a demethylase for 6mA in *Mus musculus* (Wu et al., 2016). To validate the hypothesis, we investigated whether ALKBH1 could demethylate 6mA from human genome DNA *in vitro* and *in vivo* by dot blotting and LC-MS/MS assays. We found that silencing of *ALKBH1* increased the 6mA modification level in human genome DNA (Fig. 4A), while ectopic expression of *ALKBH1* decreased the 6mA modification level in a dose-dependent manner (Fig. 4B). However, silencing or over-expression of *ALKBH1* did not change RNA m6A levels (see also Fig. S6E-S6H).

**Figure 4.**
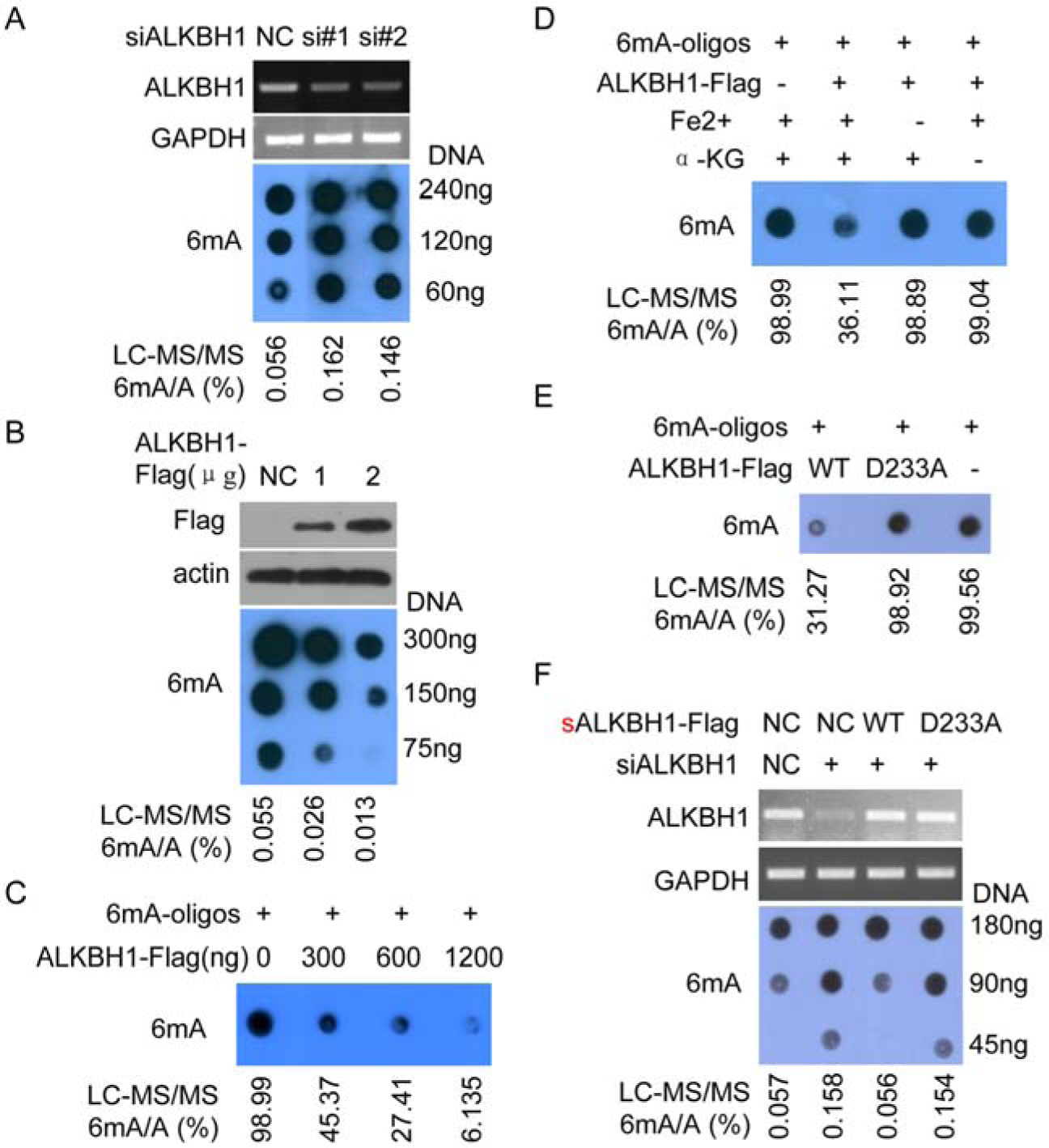
ALKBH1 is a demethylase for DNA 6mA demethylation in human, and D233 is the catalytic site of ALKBH1. (A) BEL-7402 cells were transfected with anti-ALKBH1 siRNAs, then the ALKBH1 levels were detected by RT-PCR (upper panel), and the 6mA levels of genomic DNA were subjected to dot blotting (middle panel) and LC-MS/MS (low panel) assays. (B) MHCC-LM3 cells were transfected with the indicated amounts of ALKBH1-Flag plasmids, then the ALKBH1 levels were determined by western blotting, and the 6mA levels of genomic DNA were subjected to dot blotting and LC-MS/MS assays. (C) Recombinant ALKBH1 directly demethylated the 6mA modification in the 6mA-DNA oligos in vitro demethylation reaction. The 6mA levels were detected by dot blotting (upper panel) and LC-MS/MS (low panel) assays. (D) ALKBH1 demethylase activities were dependent on Fe^2+^ and α-KG in vitro demethylation reaction. (E) Recombinant ALKBH1 D233A mutant did not demethylate the DNA oligos compared to recombinant wild type ALKBH1 in the *in vitro* methylation reaction. (F) SK-hep1 cells were co-transfected with anti-ALKBH1 siRNAs together with the indicated synonymous mutant sALKBH1-Flag plasmids, the ALKBH1 levels were determined by RT-PCR (upper panel), the 6mA levels were determined by dot blotting (middle panel) and LC-MS/MS (low panel) assays. See also Figure S5 and S6.

Next, we performed an *in vitro* demethylation assay to further determine whether ALKBH1 is a direct 6mA demethylase in human. We showed that the recombinant ALKBH1-Flag could directly and efficiently decrease 6mA modification level from the synthetic 6mA-modificated oligonucleotide substrates in a dose-dependent manner (Fig. 4C). We further demonstrated that the demethylation activities of ALKBH1 were dependent on Fe2^+^ and 2-oxoglutarate (Fig. 4D), as expected for an active dioxygenase.

The demethylase ALKBH1 for DNA 6mA demethylation in human was highly homologous with the reported demethylase ALKBH1 in mouse. The catalytic activities of mouse ALKBH1 depend on its critical residue D233 which coordinates the Fe^2+^ ion. Therefore, the D233 in human ALKBH1 was mutated to D233A. The recombinant ALKBH1 D233A mutant did not could directly and efficiently decrease 6mA modification level from the synthetic 6mA-modificated oligonucleotide substrates in an *in vitro* demethylation assay, compared to wild-type ALKBH1 (Fig. 4E). The increase of genomics 6mA induced by silencing of *ALKBH1* in cancer cells could be efficiently rescued by ectopic expression of wild-type but not D233A mutant *sALKBH1* in which the *ALKBH1* sequences targeted by anti-*ALKBH1* siRNA were synonymously mutated (Fig. 4F). Furthermore, we generated ALKBH1 KO HEK293T cell line HEK293T-ALKBH1^null^ via CRISPR-Cas9 technology (See also Fig. S7A and S7B). Dot blotting and LC-MS/MS analyses showed that genomics DNA 6mA levels were markedly increased in ALKBH1 knockout HEK293T cells (See also lane 1 and 2 in Fig. S7C). Ectopic expression of *ALKBH1* attenuated the enhancement of genomics DNA 6mA levels in HEK293T-ALKBH1^null^ cells, whereas the *ALKBH1* D233A mutant did not (See also Fig. S7C). Collectively, our results indicated that ALKBH1 is a demethylase responsible for DNA 6mA modification in the human genome.

### Low genomic 6mA level indicates a poor prognosis for human cancer patients

To further investigate the functional significance of the DNA 6mA modification, the genomic 6mA levels were further analyzed in human cancers and their corresponding adjacent non-tumoral tissues by dot blotting and LC-MS/MS assays. The genomic 6mA and its methyltransferase N6AMT1 levels were frequently down-regulated in these primary cancer tissues (T) compared with those in their corresponding non-tumoral (N) tissues, while the 6mA demethylase ALKBH1 levels were frequently up-regulated in these primary cancer tissues (Fig. 5A and 5B). The microarray gene expression data deposited in the Oncomine database also demonstrated that *N6AMT1* mRNA level was down-regulated and *ALKBH1* was up-regulated in human gastric and liver cancer tissues compared to normal tissues (See also Fig. S8A and S8C). We further found that *N6AMT1* gene copy number was decreased in gastric and liver cancer tissues (See also Fig. S8B), but *ALKBH1* was not (See also Fig. S8D), compared to normal tissues, suggesting that *N6AMT1* down-regulation could be induced by *N6AMT1* gene loss in cancer cells. The genomics 6mA levels were positively and negatively correlated with *N6AMT1* and *ALKBH1* mRNA levels in human tissues, respectively (Fig. 5C).

**Figure 5.**
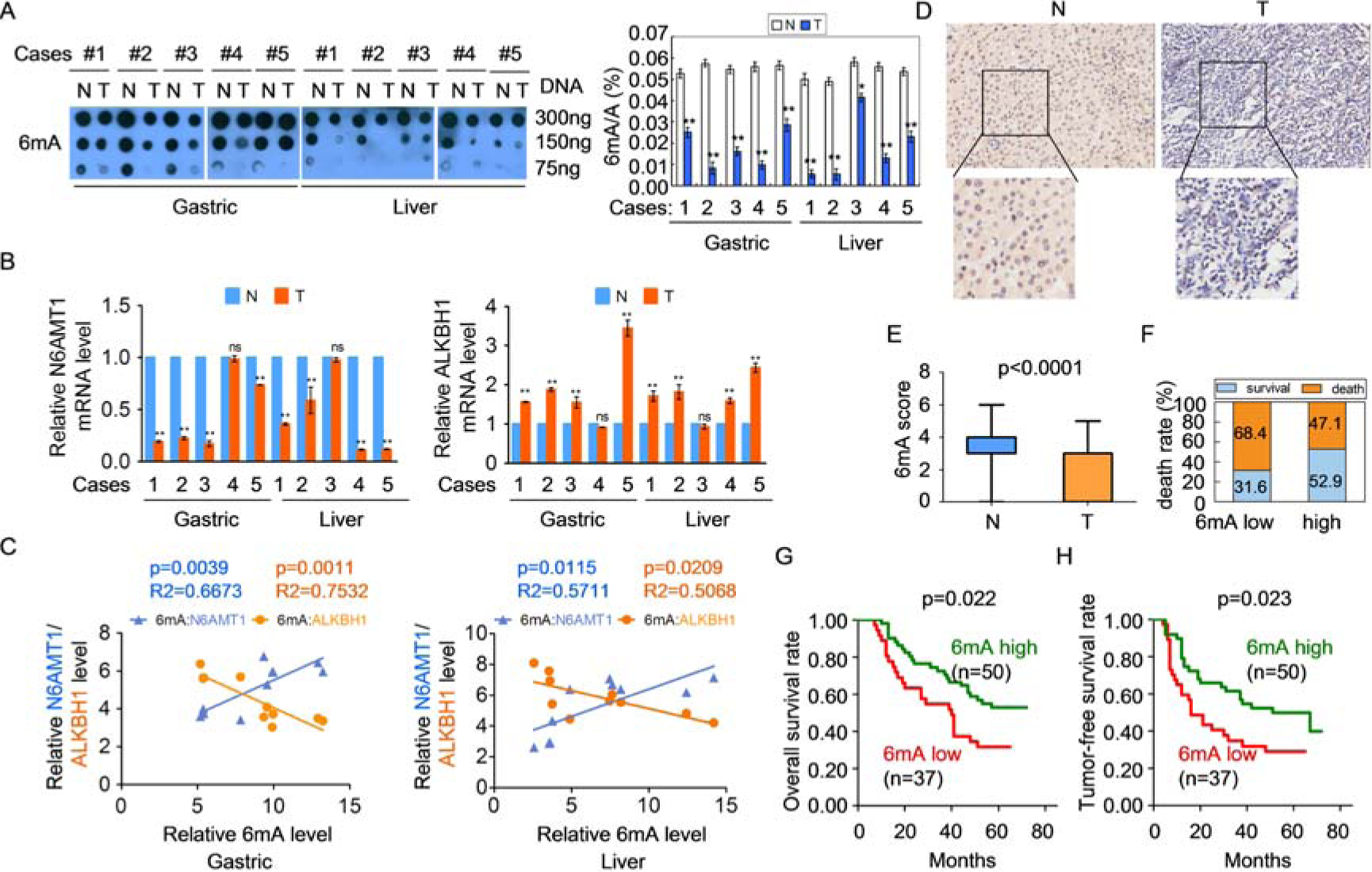
Low genomic DNA 6mA level is associated with a poor prognosis in cancer patients. (A) The genomic DNA 6mA levels were detected by dot blotting in ten pairs of matched primary gastric and liver cancer tissues (T) and corresponding adjacent non-tumoral gastric and liver tissues (N) by dot blotting (left panel) and LC-MS/MS (right panel) assays. (B) The mRNA levels of *N6AMT1* and *ALKBH1* in human tissues as (A) were detected by real-time PCR. (C) The genomic DNA 6mA levels were positively and negatively correlated with the *N6AMT1* and *ALKBH1* mRNA levels in human cancer tissues, respectively. (D) The genomic DNA 6mA levels in 87 pairs of matched liver cancer tissues (T) and their corresponding N tissues were detected by IHC assay with an anti-6mA antibody. Representative IHC images of genomic DNA 6mA level in T and their corresponding N tissues. (E) Differences in genomics 6mA level scores between T and the corresponding N tissues in Figure 5D are presented as a box plot (n=87). (F) Associations between genomic 6mA levels and the percentage of patient death were analyzed in liver cancer samples. (G, H) Kaplan-Meier plots for the overall patient survival rate (G) and the tumor-free survival rate (H) of patients with liver cancer in genomic 6mA low and 6mA high groups from Figure 5D (Log rank test). The data are represented as the means ± SEM. See also Figure S8 and Table S6.

Furthermore, an extensive tissue microarray analysis of 87 pairs of matched liver cancer (T) and corresponding adjacent non-tumor liver tissue (N) samples was performed to investigate the genomic 6mA levels using an IHC assay with an anti-6mA specific antibody (Fig. 5D). Reduced genomic 6mA levels were detected in liver cancer (T) tissues compared with those in N tissues (Fig. 5E). Reduced genomic 6mA levels in liver cancer tissues were positively associated with tumor size, histological grade, AFP level, tumor recurrence and TNM stage of liver cancer (See also Table S6). Higher patient death rates were observed in liver cancer patients with genomic 6mA low level compared with the patients classified as genomics 6mA high level (Fig. 5F). Kaplan-Meier survival analyses revealed that patients with lower genomic 6mA levels were at increased risk of cancer-related death compared with patients with higher genomic 6mA levels (Fig. 5G and 5H, log-rank test). Collectively, our results indicated that reduced genomic 6mA levels were correlated with a poor prognosis for human cancer patients, and the DNA 6mA modification may be involved in the human disease processes such as carcinogenesis.

### Decrease of genomic DNA 6mA levels promotes tumorigenesis

To further investigate the effects of DNA 6mA modification on tumorigenesis, *N6AMT1* and *ALKBH1* were over-expressed and silenced in cancer cells. We found that silencing of *N6AMT1*, which decreased cellular genomics DNA 6mA level, promoted cancer cell growth, colony formation, migration and invasion (Fig. 6A-6C, see also Fig. S9). In contrast, *N6AMT1* over-expression inhibited cancer cell growth, colony formation, migration and invasion in a dose-dependent manner (See also Fig. S10). Silencing of *ALKBH1*, which increased cellular genomic DNA 6mA levels, inhibited cancer cell growth, colony formation, migration and invasion (Fig. 7A-7C, see also Fig. S11), while *ALKBH1* over-expression reversed these effects in a dose-dependent manner in cancer cells (See also Fig. S12).

**Figure 6.**
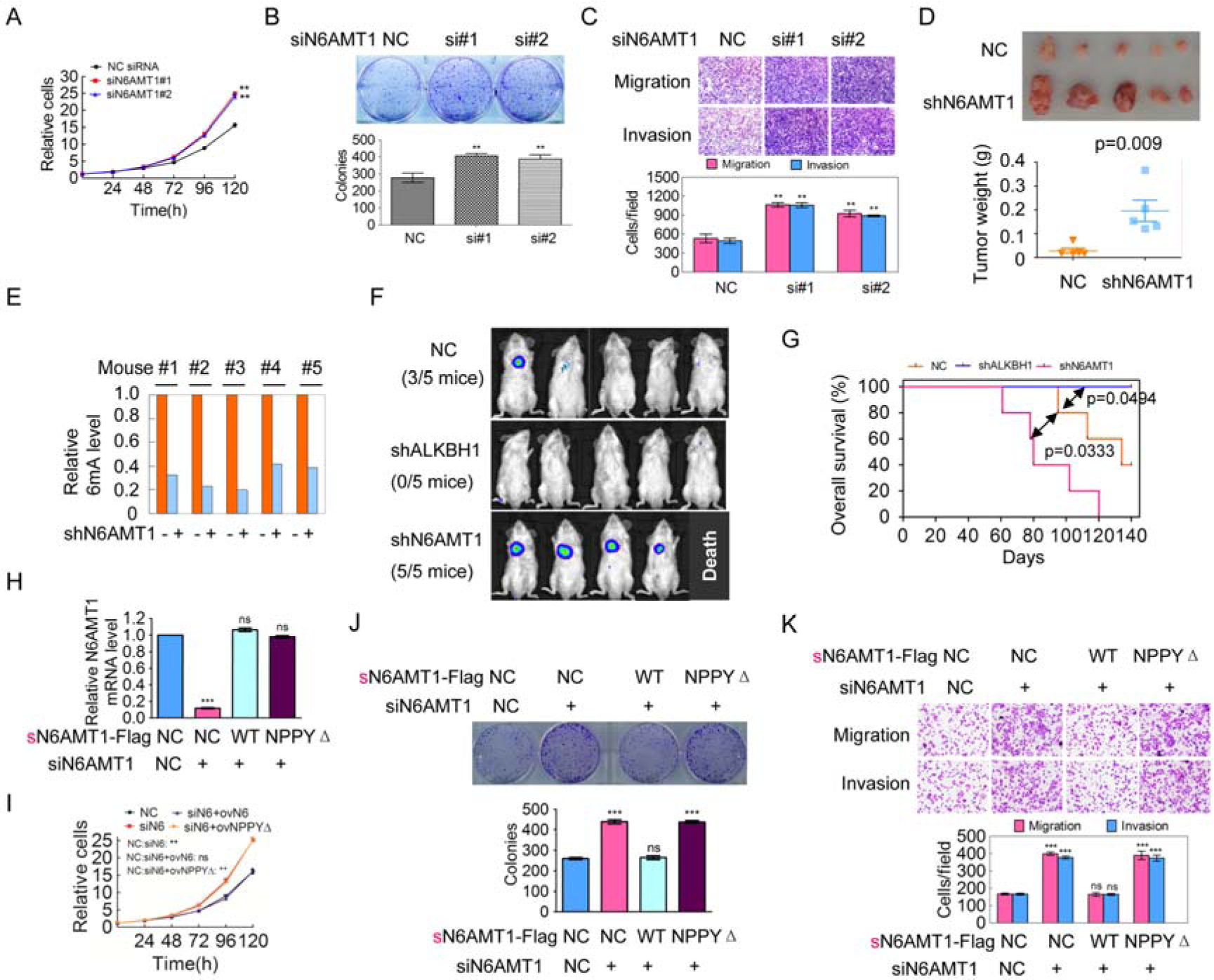
Decrease of genomic 6mA modification level promotes tumorigenesis in cancer cells. (A-C) SK-hep1 cells were transfected with anti-N6AMT1 siRNAs, the cell growth (A), colony formation (upper panel in B), migration and invasion (upper panel in C) abilities were determined. The colony number (low panel in B) and migrated and invasive cancer cell number (low panel in C) were counted (n=3). (D) The *in vivo* growth of the cancer cells stably silencing N6AMT1 was examined. Mouse xenograft tumors are shown in the upper panel. The weights of the xenograft tumors are presented in the low sides (n=5). (E) The 6mA levels in mouse xenograft tumors were determined by dot blotting, and the densities of each dot were quantified and normalized. (F) The *in vivo* metastasis of the cancer cells stably silencing N6AMT1 or ALKBH1 was examined. NOD-SCID mice were transplanted with the indicated Luc-labeled SK-hep1 cells (2×10^6^ cells/mouse) via tail vein injection; luciferase activity was visualized nine weeks post-transplantation (n=5). (G) Kaplan-Meier curves are shown for three cohorts of transplanted mice carrying SK-hep1-Luc-NC shRNA, SK-hep1-Luc-N6AMT1 shRNA and SK-hep1-Luc-ALKBH1 shRNA cells (n=5). (H-K) SK-hep1 cells were co-transfected with anti-N6AMT1 siRNAs together with the indicated synonymous mutant sN6AMT1-Flag plasmids, the N6AMT1 levels were determined by real-time PCR (H), the cell growth (I), colony formation (J), migration and invasion (K) abilities were determined. The data are represented as the means ± SEM. See also Figure S9 and S10.

**Figure 7.**
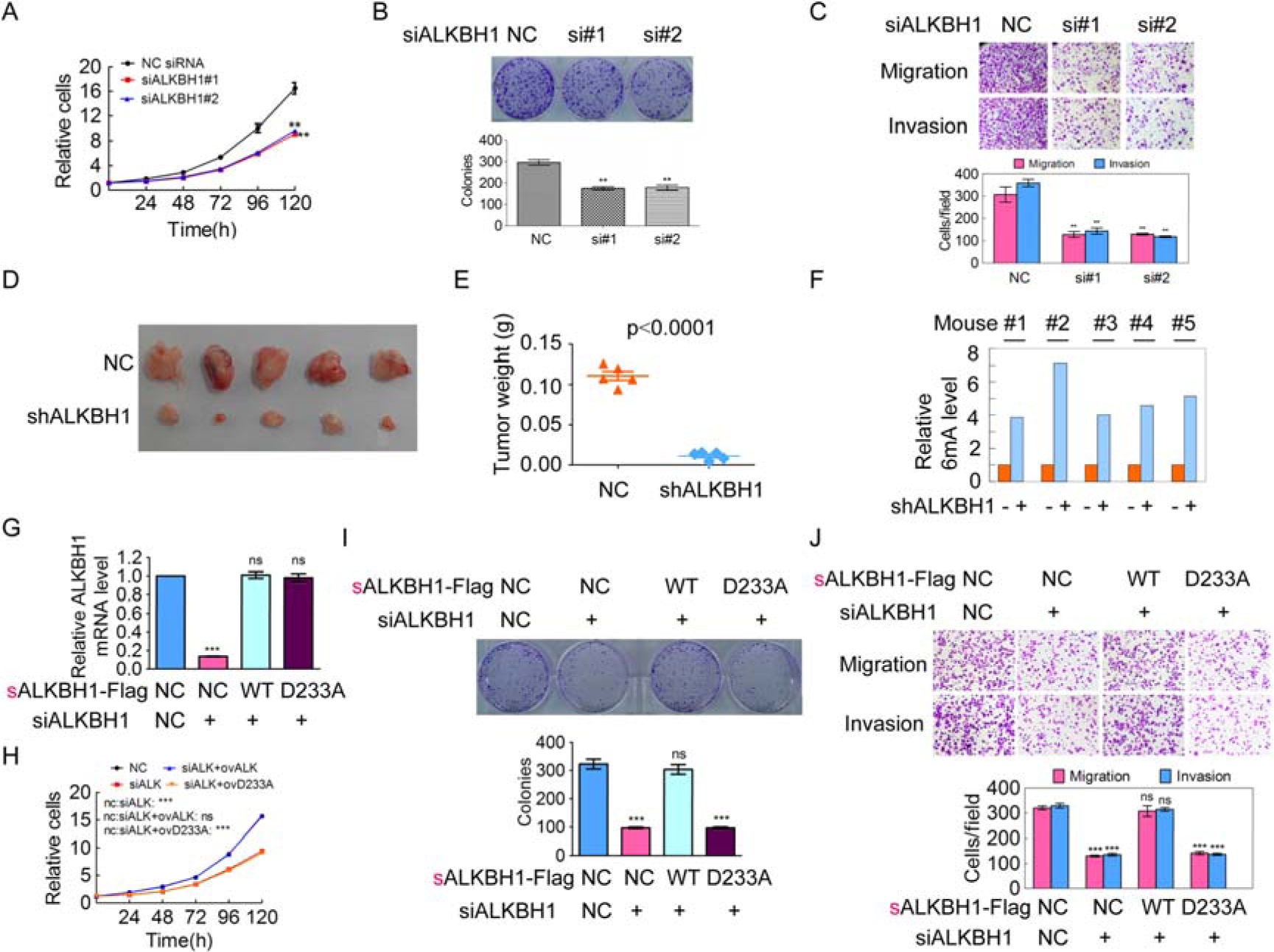
Increase of genomic 6mA modification level inhibits tumorigenesis in cancer cells. (A-C) SMMC-7721 cells were transfected with anti-ALKBH1 siRNAs, the cell growth (A), colony formation (B), migration and invasion (C) abilities were determined (n=3). (D) The *in vivo* growth of the cancer cells stably silencing ALKBH1 was examined. (E) The weights of the xenograft tumors presented in Figure 7D are analyzed (n=5). (F) The 6mA levels in mouse xenograft tumors were determined by dot blotting, and the densities of each dot were quantified and normalized. (G-J) SK-hep1 cells were co-transfected with anti-ALKBH1 siRNAs together with the indicated synonymous mutant sALKBH1-Flag plasmids, the ALKBH1 levels were determined by real-time PCR (G), the cell growth (H), colony formation (I), migration and invasion (J) abilities were determined. The data are represented as the means ± SEM. See also Figure S11 and S12.

More importantly, as shown in Figure 6D, the *in vivo* growth of cancer xenografts composed of N6AMT1 expression-stably silenced cells was significantly increased. The 6mA levels in mouse xenograft cancer tissues which are composed of N6AMT1 expression-stably silenced cells were decreased compared to those which are composed of the negative control cancer cells (Fig 6E). However, the *in vivo* growth of cancer xenografts composed of ALKBH1 expression-stably silenced cells was significantly impaired (Fig. 7D and 7E). The 6mA levels in mouse xenograft cancer tissues with stable ALKBH1 knockdown were increased compared to those in negative control (Fig. 7F).

Furthermore, larger metastatic nodules developed in mouse lungs after tail vein injections with luciferase-tagged SK-hep1 cells in which the expression of *N6AMT1* was stably silenced, smaller metastatic or non-metastastatic nodules developed in mouse lungs with SK-hep1 cells in which the expression of *ALKBH1* was stably silenced, compared to those with negative control SK-hep1 cells (Fig. 6F). Kaplan-Meier survival analyses revealed that mice with cancer xenografts carrying *N6AMT1*-stable silencing cells were at increased risk of cancer-related death compared with mice with cancer xenografts carrying NC cells (p=0.033, log-rank test), while mice with cancer xenografts carrying *ALKBH1*-stable silencing cells were at decreased risk of cancer-related death compared with mice with cancer xenografts carrying NC cells (p=0.049, log-rank test).

We furthermore demonstrated that the increase of cell growth, colony formation, migration and invasion abilities induced by silencing of *N6AMT1* in cancer cells could be efficiently rescued by ectopic expression of wild-type but not NPPYΔ mutant *sN6AMT1* in which the *N6AMT1* sequences targeted by anti-*N6AMT1* siRNA were synonymously mutated so that *sN6AMT1* and *sN6AMT1* NPPYΔ mutant expression was not silenced by anti-*N6AMT1* siRNA (Fig. 6H-6K). And the reduction of cell growth, colony formation, migration and invasion abilities induced by silencing of ALKBH1 in cancer cells could be efficiently rescued by ectopic expression of wild-type but not D233A mutant *sALKBH1* in which the *ALKBH1* sequences targeted by anti-*ALKBH1* siRNA were synonymously mutated so that *sALKBH1* and *sALKBH1* D233A mutant expression was not silenced by anti-*ALKBH1* siRNA (Fig. 7G-7J). Collectively, our results suggested that decrease of genomic DNA 6mA modification levels in human cancer cells can promote tumorigenesis.

## Discussion

In this study, we found that DNA 6mA was extensively present in the human genome, especially in the mitochondria and that [G/C]AGG[C/T] was the most prevalent motif for 6mA modification. In addition, 6mA density was enriched in the exon regions and was associated with actively transcribed genes in the human cells. We identified methyltransferase N6AMT1 and demethylase ALKBH1 as responsible for the DNA 6mA modification in human. Interestingly, we performed exploratory analysis and demonstrated lower 6mA level in human cancer tissues compared to adjacent non-tumoral tissues, and that the decrease of genomic DNA 6mA levels promotes human tumorigenesis.

We compiled a catalog of all DNA 6mA modification sites in blood lymphocyte samples in this study. Our conclusion that 6mA is extensively present in human genome DNA was supported by multiple lines of evidence. First, 6mA on human genome DNA was detected using SMRT sequencing (103X genome-wide coverage). Although SMRT sequencing has difficulty distinguishing between 6mA and 1mA, our LC-MS/MS assay demonstrated that the DNA sample subjected to SMRT sequencing had undetectable 1mA, indicating that our identified DNA methylation modifications likely represent 6mA. Second, the DNA 6mA modification identified by SMRT sequencing was validated by 6mA-IP-seq. The 6mA-containing DNA regions identified by 6mA-IP-seq were highly overlapped with the 6mA site-occupied regions identified by SMRT sequencing. We demonstrated similar 6mA distribution patterns, motif and correlation with gene transcription activation between SMRT sequencing and 6mA-IP-seq, despite differences in sensitivity of the technologies. Third, the presence of 6mA was identified by LC-MS/MS assay. The 6mA density identified by SMRT sequencing (0.051%) and LC-MS/MS (0.056%) was similar. Fourth, the presence of 6mA was also identified by 6mA-specific antibody assays (including 6mA-IP-qPCR and dot blotting). Fifth, we identified the methyltransferase N6AMT1 and demethylase ALKBH1 for addition and removal of 6mA in human genome DNA, not m6A in RNA. Sixth, the catalytic motif NPPY at N6AMT1 and the critical residue D233 at ALKBH1 for addition and removal of 6mA were identified. Finally, the 6mA level was down-regulated in human cancer tissues, its level was positively correlated with N6AMT1 and negatively correlated with ALKBH1, and the reduction of genomics 6mA promoted tumorigenesis.

Our SMRT sequencing data identified a broad 6mA genomic distribution with the most prevalent motif sequence being [G/C]AGG[C/T] in human genome DNA, similar to one of two motifs (GAGG and AGAA) in the *C. elegans* genome (Greer et al., 2015), but different from those in *Chlamydomonas* (AT in TSS region) (Fu et al., 2015), in mouse (GAA, G[A/G]AATA)(Wu et al., 2016), and significantly different from that in prokaryotes in which most of the 6mA sites are located with the palindromic sequence (Touzain et al., 2011). These observations indicated that the 6mA modification pattern was not conserved across the eukaryotes specie and was species-specific, likely reflecting the potential diverse biological functions and mechanisms. Recent studies from a limited number of eukaryotes have exhibited the differences in function of 6mA. Our and other’s studies indicated that DNA 6mA modification marks activated gene transcription in human, C. *elegans (Greer et al., 2015)*, C. *reinhardtii* (Fu et al., 2015), and fungi (Mondo et al., 2017), and transposon expression in *D. melanogaster* (Zhang et al., 2015), while suppresses gene expression in mouse (Wu et al., 2016). We further found that the 6mA-enriched regions in genome that were associated with the activation of gene expression were also different in the different eukaryotes. 6mA was enriched in the exon regions in human, but enriched in TSS in C. *reinhardtii (Fu et al., 2015)*, or transposons in *D. melanogaster* (Zhang et al., 2015). The detailed regulatory mechanism of 6mA on gene and transposon expression is still unclear and deserves further investigation in future studies.

Searching for specific 6mA methyltransferase and demethylase is important for understanding the potential role of 6mA in human cells. In this study, we identified methyltransferase N6AMT1 and demethylase ALKBH1 as responsible for the addition and removal of 6mA in human genomic DNA, respectively. The demethylase ALKBH1 for DNA 6mA demethylation in human was highly homologous with the reported demethylase ALKBH1 in mouse, but was different from the demethylase DMAD in *Drosophila* (Zhang et al., 2015), NMAD-1 in C.*elegans (Greer et al., 2015)*. And the catalytic critical residue D233 in human ALKBH1 is the same with that in mouse ALKBH1. However, both NMAD-1 and ALKBH1 belong to AlkB family proteins which contain the conserved Fe2+ ion and 2-oxo-glutarate-dependent, dioxygenase domain (Liu et al., 2016a; Westbye et al., 2008). Additionally, the AlkB family proteins, such as ALKBH5 and FTO in mammals, also catalyze the demethylation of 6mA in RNA (Jia et al., 2011; Zheng et al., 2013). Taken together, these data suggested that the demethylases for DNA 6mA demethylation may be evolutionarily conserved in mammals and belong to the AlkB family.

The methyltransferase for 6mA addition in genomic DNA was not discovered in previous report on fungi (Mondo et al., 2017), *C. reinhardtii* (Fu et al., 2015), *D. melanogaster* (Zhang et al., 2015) and mouse, except from C. *elegans* (Greer et al., 2015). Here, we identify N6AMT1 as a DNA 6mA methyltransferase in human. However, our identified methyltransferase N6AMT1 was not homologous with the methyltransferase DAMT-1 in C.*elegans*. The DNA 6mA methylase DAMT-1 in *C. elegans* is a member of MT-A70 domain family evolving from the m.MunI-like bacterial 6mA DNA methyltransferase. But, in human, METTL3, which belongs to MT-A70 family, is a RNA m6A methyltransferase, not a DNA 6mA methyltransferase (Liu et al., 2014). Interestingly and surprisingly, our identified human DNA 6mA methyltransferase N6AMT1 has regions of sequence similarity with M.TaqI-like 6mA DNA methyltransferases of bacterial restriction-modification systems (Timinskas et al., 1995). Both of them contain an adenine methyltransferase-characterized amino acid motif (D/N/S) PP (Y/F/W). The catalytic motif NPPY in human N6AMT1 is critical for the catalytic activities of N6AMT1, suggesting that human DNA 6mA methyltransferase may share an evolutionary origin with bacteria.

Importantly and interestingly, the 6mA modification level was down-regulated in human cancer tissues compared to their adjacent non-tumoral tissues, accompanying *N6AMT1* down-regulation and gene copy number loss and *ALKBH1* up-regulation. The significantly positive and negative associations of the 6mA level with the *N6AMT1* and *ALKBH1* expression level were observed in human cancer tissues, respectively. Over-expression of *N6AMT1* and silencing of *ALKBH1*, which increased genomic DNA 6mA levels, inhibited cancer cell growth, colony formation, migration and invasion *in vitro*, and tumorigenesis and metastasis *in vivo*, while over-expression of ALKBH1 and silencing of N6AMT1, which decreased genomic DNA 6mA level, exhibited oncogenic effects. These data indicated that genomics DNA 6mA modification plays crucial role in human diseases such as cancer. The DNA 5mC methylation biomarkers in genomic DNA including circulating tumor DNA (ctDNA) in liquid biopsy have been used in cancer diagnosis, detection, classification and prognosis (Costa-Pinheiro et al., 2015; Ma et al., 2015). Reduced DNA 6mA level was correlated with malignant phenotype and poor prognosis for cancer patients. The DNA 6mA methylation modification may also be a potential biomarker for cancer diagnosis and therapy.

In summary, we found that DNA 6mA modification is extensively present in human genome DNA, and [G/C]AGG[C/T] is the most prevalent motif of the 6mA modification. The 6mA modification is associated with actively transcribed genes. N6AMT1 and ALKBH1 were identified as the methyltransferase and demethylase for genomics DNA 6mA modification in human, respectively. The 6mA level is down-regulated in human cancer tissues, accompanying *N6AMT1* down-regulation and *ALKNH1* up-regulation, while the decrease of genomic DNA 6mA level promotes tumorigenesis. The discovery of 6mA in human cells sheds new light on epigenetic regulation during human diseases such as cancer.

## STAR Methods

### Tissue and DNA samples

Matched primary cancer tissues and their corresponding adjacent non-tumoral tissues were collected from gastric and liver cancer patients at the Third Affiliated Hospital of Guangzhou Medical University and Sun Yat-sen University Cancer Center (Xu et al., 2015). These cancer patients had not received preoperative anticancer treatment. Informed consent was obtained from each patient, and the collection of tissue specimens was approved by the Internal Review and Ethics Boards at the Third Affiliated Hospital of Guangzhou Medical University. Tissue microarray chips containing 90 pairs of liver cancer tissue samples matched to their adjacent non-tumoral liver tissue samples and the associated clinicopathological information were purchased from Shanghai OUTDO Biotech Co., Ltd. (Shanghai).

Blood genomics DNA samples used in the validation of 6mA modification (Measurement of the 6mA/A ratio and 6mA-IP-qPCR) were from our group-preserved samples which were previously used in long-read DNA SMRT sequencing and short-read RNA sequencing (SRX1424851 and SRX1423750 in NCBI project PRJNA301527) (Shi et al., 2016).

### Identification of 6mA modifications in human genomic DNA

The raw h5 format files of PacBio sequencing reads were downloaded from the NCBI SRA database (SRX1424851 in PRJNA301527) (Shi et al., 2016). We used the PacBio SMRT analysis platform (version 2.3.0) for DNA 6mA modification detection (http://www.pacb.com/products-and-services/analytical-software/smrt-analysis/analysis-applications/epigenetics/). Briefly, DNA raw data were aligned to the hg38 reference genome by pbalign. The polymerase kinetics information was further loaded after alignment by loadChemistry.py and loadPulses scripts from raw h5 format files. The two post-aligned datasets were then merged and sorted by using cmph5tools. The 6mA modification was identified by using ipdSummary.py script. We further filtered 6mAs with less than 25X coverage in autosome chromosomes and 12X in sex chromosomes. High-quality 6mAs were annotated by ANNOVAR (Wang et al., 2010) based on RefGene annotation file.

### 6mA-IP-seq

Genomic DNA was extracted from human blood of the HX1 sample, treated with RNase A and sonicated to produce fragments of 200-400 bp. Fragmented DNA was incubated for 2 h at 4°C with anti-6mA polyclonal antibody (Synaptic Systems) in immunoprecipitation buffer (10 mM sodium phosphate, 2 mM potassium phosphate, 137 mM NaCl, 2.7 mM KCl, 0.05% Triton X-100, pH 7.4). The mixture was then immunoprecipitated by incubation with Protein A/G Plus-Agarose (Santa Cruz) that was pre-bound with bovine serum albumin (BSA) at 4 °C for additional 2 h. After extensive washing, bound DNA was eluted from the beads in elution buffer (50 mM Tris, pH 8.0, 1% SDS, 10 mM EDTA) at 65 °C, treated with proteinase K and purified using QIAquick PCR Purification Kit (Qiagen). Four libraries for two biological replicates each for input and immunoprecipitated DNA were prepared with NEBNext DNA Library Prep Kit (NEB) according to the manufacturer’s instructions, and sequenced on the Illumina HiSeq X10 platform with 150-bp paired-end.

Raw data were trimmed with Skewer (Jiang et al., 2014) to remove adapters and low-quality bases. Reads with more than 10% N or more than 50% low-quality base were filtered out. BWA (Li and Durbin, 2009) was used to align reads to the NCBI human reference genome (hg19). After alignment, 6mA enriched regions were called with MACS version 2 (Zhang et al., 2008) by default parameters. Sequencing data have been deposited into the Gene Expression Omnibus (GEO) under the accession number GSE 104475.

### Measurement of the 6mA/A ratio by LC-MS/MS

Blood genomic DNA, RNA and synthetic oligonucleotide substrates was digested into single nucleosides in a digestion buffer containing phosphodiesterase I (0.01 U), nuclease P1 (1 U), 2mM zinc chloride and 30mM sodium acetate at pH 6.8 for 3h at 37°C, and dephosphorylated with bacterial alkaline phosphatase (10 U) for 1h at 37°C. Enzymes were removed by filtration (Amicon Ultra 10K MWCO). Individual nucleosides were resolved on a Hypersil GOLD aQ reverse phase column (Thermo Scientific), and samples were then subjected to liquid chromatography coupled with tandem mass spectrometry (LC-MS/MS) analysis on an Agilent 6490 Triple Quadrupole mass spectrometer. Nucleosides were quantified using the nucleoside-to-base ion mass transitions of 266.1 to 150.1 for DNA 6mA and 1mA, 252.1 to 136.1 for A, 282.1-150.1 for RNA m6A.

### 6mA-IP-qPCR

6mA-IP-qPCR was performed using freshly drawn blood samples harvested independently of the blood samples for SMRT sequencing. Blood genomic DNA was extracted from freshly drawn blood samples and treated with RNase to remove RNA contamination. The DNA sample was sonicated to produce ∼250bp fragments. The fragmented blood DNA was incubated with specific anti-6mA antibody (Synaptic Systems) in immunoprecipitation buffer (2.7 mM KCl, 2 mM potassium phosphate, 137 mM NaCl, 10 mM sodium phosphate, 0.05% Triton X-100, pH 7.4) for 2h at 4°C. The mixture was then immunoprecipitated by Protein A/G Plus-Agarose (Santa Cruz) that was pre-bound with bovine serum albumin (BSA) at 4°C for 2h. After extensive washing, the bound DNA was eluted from the beads in elution buffer (50 mM Tris, pH 8.0, 1% SDS, 10 mM EDTA) at 65°C, treated with proteinase K and purified using QIAquick PCR Purification Kit (Qiagen). Fold-enrichment of each fragment was determined by quantitative real-time PCR. The primers were listed in Supplementary Table S7.

### Bioinformatics analysis

The genome-wide 6mA profiling across all chromosomes were generated from Circos (Krzywinski et al., 2009). For each 6mA modification site, we extracted 20bp from the upstream and downstream sequences of the 6mA site. MEME program was then used to identify motifs in flank region (Bailey et al., 2009).

The short-read RNA sequencing raw data were downloaded from the NCBI database (SRX1423750) (Shi et al., 2016). RNA-seq short-read were aligned to the hg38 reference genome by HISAT (Kim et al., 2015). The Reads per Kilobase of transcript per Million mapped reads (RPKM) measured by RNA-SeQC was used for quantification of gene expression (DeLuca et al., 2012).

Web Gestalt was used to test the statistical enrichment of genes in Gene Ontology (Wang et al., 2017). The p-values were adjusted by the Benjamini and Hochberg’s approach, and the adjusted P-value (FDR) <0.01 was considered to be statistically significant.

### Dot blotting for DNA 6mA

Dot blotting was performed as previously described with minor modification (Zhang et al., 2015). Briefly, the extracted DNA was treated with RNase A and denatured at 95°C for 10min and cooled down on ice for 5min. DNA was then loaded on the nitrocellulose membrane (Protran BA 85, Whatman), air dry for 5min. The membrane was baked at 80°C for 2h and then blocked in the blocking buffer (1% BSA and 5% milk in PBST) for 1.5h at room temperature. The membranes were incubated with a specific anti-6mA antibody (Millipore, 1:1000) overnight at 4°C, followed by incubation with HRP-conjugated anti-rabbit IgG secondary antibody (Bioworld, 1:4000) at room temperature for 1.5h. The antibody-bound 6mA was detected by using the enhanced chemiluminescence Supersignal West Pico kit (Thermo, USA) followed by exposure to autoradiographic film in darkroom.

### Dot blotting for RNA m6A

RNA dot blotting was performed as previously described with minor modification (Ping et al., 2014). Briefly, the extracted total RNA was placed at 55℃ for 15min to denature in a denaturing buffer (2.2M formaldehyde, 50% deionized formamide, 0.5х MOPS buffer). RNA was heated at 95℃ for 10 min and cooled down on ice for 5min. RNA was then spotted onto Amersham Hybond-N+ membranes (GE Healthcare, cat# RPN303B), air dry for 5min. The RNA m6A detection in the membrane was the same as DNA dot blotting.

### N6AMT1 methylase assays in vitro

In vitro methylation reactions were performed as previously described with minor modification (Banerjee and Rao, 2011). Briefly, the reactions were performed in a 25μl methylation reaction buffer containing 250 pmol DNA oligos, 800ng recombinant N6AMT1-Flag protein or N6AMT1-Flag NPPYΔ mutant protein, 50mM Tris-HCl (pH =7.6), 50mM KCl, 10mM Mg(OAc)_2_, 7mM β-mercaptoethanol, 800μM S-adenosylmethionine, and 100μg/ml bovine serum albumin. Reactions were carried out overnight at 25°C and stopped by protein inactivation at 95°C for 10min. DNA was purified (DP214, Tiangen Biotech) and 2μl of purified DNA was used for 6mA dot blotting. The following four oligos were used: oligo1#: CATGATACCTTATGGA<u>A</u>AGCATGCTTGTATTTCTTATGAA CCATGATACCTTATGGAAAGCATGCTTGTATTTCTTATGAAC; oligo2#: CCGC GG<u>A</u>GGCTGCT; oligo#3: CCGCGC<u>A</u>GGCTGCT; oligi#4: CCGCGG<u>A</u>GT**C**TGCT. The underlined A in the four oligos were mutated to G and produced the A>G mutant oligos.

### ALKBH1 demethylase assays in vitro

In vitro demethylation reactions were performed as previously described with minor modification (Wu et al., 2016). Briefly, the reactions were performed in a 50μl demethylation reaction buffer containing 50pmol DNA oligos, 500ng recombinant ALKBH1-Flag protein or ALKBH1-Flag D233A mutant protein, 50µM HEPES (pH =7.0), 50µM KCl, 1mM MgCl_2_, 2mM ascorbic acid, 1mM α-KG, and 1mM (NH_4_)_2_Fe(SO_4_)_2·_6H_2_O. Reactions were carried out for 1h at 37°C and stopped with 5 mM EDTA followed by heating at 95°C for 10min. Then 2μl of a reaction product was used for dot blotting. The 6mA-oligos used here as previously described (Wu et al., 2016).

### ALKBH1 and N6AMT1 knockout cell lines by CRISPR-Cas9

The *ALKBH1* and *N6AMT1* Cas9/sgRNAs plasmids pGE-4 (pU6gRNA1Cas9puroU6gRNA2) were purchased from Genepharma (Shanghai, China). The two sgRNA targeting sites in *ALKBH1* were 5’-TCAGAGCCGGCCCGGGACCG-3’ (site1) and 5’-GGCCCACGCAGCCCGTGG CA-3’ (site2). The two sgRNA targeting sites in *N6AMT1* were 5’- CGGGCACGTGGGCCGCGGCG-3’ (site1) and 5’-CGACGTGTACGAGCCCGCG G-3’ (site2). These plasmids were transfected into HEK293T cells by Lipo2000 (Invitrogen). Two days after transfection, HEK293T cells were selected with Puromycin for two days. The cells were seeded onto 96-well plates for single-cell culture to obtain single-derived cell colonies and confirmed by PCR.

### DNA 6mA detection by Immunohistochemistry (IHC) with anti-6mA antibody

IHC assays were performed as previously described using anti-6mA antibody as previously described with minor modification (Huang et al., 2017). IHC assays were performed on tissue microarray chips with anti-6mA antibody (1:100). The tissue microarray chips were treated with 50μg/ml RNaseA for 60min prior to anti-6mA antibody incubation to remove RNA contamination. All IHC samples were assessed by two independent pathologists blinded to both the sample origins and the subject outcomes. Nuclear 6mA staining was evaluated. Briefly, each specimen was scored according to the extent of cell nuclear staining (≤10% positive cells for 0; 11-50% positive cells for 2; 51-80% positive cells for 3; >80% positive cells for 4) and the nuclear staining intensity (no staining for 0; slight staining for 1; moderate staining for 2; strong staining for 3). Scores for the percentage of positive cells and the staining intensity were added. Score of 0 was indicative of low 6mA level (low) and scores of 3-7 were indicative of high 6mA (high) in liver cancer tissues.

### Animal studies

Male BALB/c nude mice (3-4 weeks old) and NOD-SCID mice (3-4 weeks old) were purchased from Charles River Laboratories in China (Beijing). The *in vivo* tumor growth and *in vivo* metastasis assays were performed as previously described with minor modification (Huang et al., 2017). For the tumorigenesis assay, 1×10^6^ SK-hep1-Luc-NC or SK-hep1-Luc-N6AMT1 shRNA-transduced cancer cells or 1×10^6^ SK-hep1-Luc-NC or SK-hep1-Luc-ALKBH1 shRNA-transduced cancer cells were subcutaneously injected into the dorsal flanks of each mouse (n=5). After three weeks, the mice were euthanized, and the tumors were dissected and weighed. For the metastasis assay, NOD-SCID mice in each experimental group were injected with 2×10^6^ luciferase-labeled SK-hep1-Luc-NC, SK-hep1-Luc-N6AMT1 shRNA or SK-hep1-Luc-ALKBH1 shRNA-transduced cancer cells through the tail vein. Each mouse was injected with 0.15mg/g.mouse D-Luciferin Potassium Salt D in DPBS buffer (Invitrogen) for 5min through the tail vein nine weeks after implantation. The metastatic foci in the lungs were visualized using the IVIS Lumina HTX Imaging System (PerkinElmer) nine weeks after implantation.

The mice used in these experiments were bred and maintained under defined conditions at the Animal Experiment Center of the College of Medicine (SPF grade), Jinan University. The animal experiments were approved by the Laboratory Animal Ethics Committee of The Third Affiliated Hospital of Guangzhou Medicine University and Jinan University and conformed to the legal mandates and national guidelines for the care and maintenance of laboratory animals.

## Statistical analysis

Statistical analyses were performed using the Prism5 software. Two-tailed unpaired Student’s t-tests were applied for comparisons between two groups. The data are presented as the means ± SEM except where stated otherwise. The *p<0.05, **p<0.01 or ***p<0.001 values were considered statistically significant.

## Acknowledgments

This work was supported by the National Natural Science Foundation of China (81772998, 81672393), the R&D Plan of Guangzhou and Guangdong (201704020115, 2017A020215096), the Yangcheng Scholars program from the Ministry of Education of Guangzhou (1201561583), Innovative Research Team of Ministry of Education of Guangzhou (1201610015), the National Funds of Developing Local Colleges and Universities (B16056001), the Fundamental Research Funds for the Central Universities (15ykjc23d), and Guangdong Natural Science Foundation (2015A030313127), China Postdoctoral Science Foundation (2017M612798), National Institute of Health (HG006465). The authors declare competing financial interests.

## Author Contributions

G.R.Y. conceived the project and designed the experiments; C.L.X. guided informatics analyses; G.R.Y. and G.Y. performed the identification of 6mA; S.Z., G.R.Y., D.C. and J.B.L. performed the experiments for identifying for 6mA modification, detecting 6mA in samples and investigating the functions of 6mA *in vitro* and *in vivo*; X.F.G., Q.Z. and Z.L. performed LC-MS/MS and 6mA-IP-qPCR assays and 6mA-IP-seq; G.R.Y. and D.C. performed the IHC assay; M.H., Y.C., S.Q.X., F.L. and X.C.B. performed the informatics analysis; K.W. guided SMRT data analysis, advised the study and revised the manuscript; K.W., G.Y. and D.P.W. generated SMRT sequencing data, RNA-Seq data and additional samples for experimental validation; G.R.Y. collected the cancer samples and wrote the paper.

## References

1. Aas, P.A., Otterlei, M., Falnes, P.O., Vagbo, C.B., Skorpen, F., Akbari, M., Sundheim, O., Bjoras, M., Slupphaug, G., Seeberg, E., et al. (2003). Human and bacterial oxidative demethylases repair alkylation damage in both RNA and DNA. Nature 421, 859–863.

2. Bailey, T.L., Boden, M., Buske, F.A., Frith, M., Grant, C.E., Clementi, L., Ren, J., Li, W.W., and Noble, W.S. (2009). MEME SUITE: tools for motif discovery and searching. Nucleic Acids Res 37, W202–208.

3. Banerjee, A., and Rao, D.N. (2011). Functional analysis of an acid adaptive DNA adenine methyltransferase from Helicobacter pylori 26695. PLoS One 6, e16810.

4. Bergman, Y., and Cedar, H. (2013). DNA methylation dynamics in health and disease. Nat Struct Mol Biol 20, 274–281.

5. Costa-Pinheiro, P., Montezuma, D., Henrique, R., and Jeronimo, C. (2015). Diagnostic and prognostic epigenetic biomarkers in cancer. Epigenomics 7, 1003–1015.

6. DeLuca, D.S., Levin, J.Z., Sivachenko, A., Fennell, T., Nazaire, M.D., Williams, C., Reich, M., Winckler, W., and Getz, G. (2012). RNA-SeQC: RNA-seq metrics for quality control and process optimization. Bioinformatics 28, 1530–1532.

7. Duncan, T., Trewick, S.C., Koivisto, P., Bates, P.A., Lindahl, T., and Sedgwick, B. (2002). Reversal of DNA alkylation damage by two human dioxygenases. Proc Natl Acad Sci U S A 99, 16660–16665.

8. Flusberg, B.A., Webster, D.R., Lee, J.H., Travers, K.J., Olivares, E.C., Clark, T.A., Korlach, J., and Turner, S.W. (2010). Direct detection of DNA methylation during single-molecule, real-time sequencing. Nat Methods 7, 461–465.

9. Fu, Y., Luo, G.Z., Chen, K., Deng, X., Yu, M., Han, D., Hao, Z., Liu, J., Lu, X., Dore, L.C., et al. (2015). N6-methyldeoxyadenosine marks active transcription start sites in Chlamydomonas. Cell 161, 879–892.

10. Greer, E.L., Blanco, M.A., Gu, L., Sendinc, E., Liu, J., Aristizabal-Corrales, D., Hsu, C.H., Aravind, L., He, C., and Shi, Y. (2015). DNA Methylation on N6-Adenine in C. elegans. Cell 161, 868–878.

11. Huang, J.Z., Chen, M., Chen, Gao, X.C., Zhu, S., Huang, H., Hu, M., Zhu, H., and Yan, G.R. (2017). A Peptide Encoded by a Putative lncRNA HOXB-AS3 Suppresses Colon Cancer Growth. Mol Cell 68, 171–184 e176.

12. Jia, G., Fu, Y., Zhao, X., Dai, Q., Zheng, G., Yang, Y., Yi, C., Lindahl, T., Pan, T., Yang, Y.G., et al. (2011). N6-methyladenosine in nuclear RNA is a major substrate of the obesity-associated FTO. Nat Chem Biol 7, 885–887.

13. Jiang, H., Lei, R., Ding, S.W., and Zhu, S. (2014). Skewer: a fast and accurate adapter trimmer for next-generation sequencing paired-end reads. BMC Bioinformatics 15, 182.

14. Jones, P.A. (2012). Functions of DNA methylation: islands, start sites, gene bodies and beyond. Nat Rev Genet 13, 484–492.

15. Kim, D., Langmead, B., and Salzberg, S.L. (2015). HISAT: a fast spliced aligner with low memory requirements. Nat Methods 12, 357–360.

16. Krzywinski, M., Schein, J., Birol, I., Connors, J., Gascoyne, R., Horsman, D., Jones, S.J., and Marra, M.A. (2009). Circos: an information aesthetic for comparative genomics. Genome Res 19, 1639–1645.

17. Li, H., and Durbin, R. (2009). Fast and accurate short read alignment with Burrows-Wheeler transform. Bioinformatics 25, 1754–1760.

18. Liu, F., Clark, W., Luo, G., Wang, X., Fu, Y., Wei, J., Wang, X., Hao, Z., Dai, Q., Zheng, G., et al. (2016a). ALKBH1-Mediated tRNA Demethylation Regulates Translation. Cell 167, 816–828 e816.

19. Liu, J., Yue, Y., Han, D., Wang, X., Fu, Y., Zhang, L., Jia, G., Yu, M., Lu, Z., Deng, X., et al. (2014). A METTL3-METTL14 complex mediates mammalian nuclear RNA N6-adenosine methylation. Nat Chem Biol 10, 93–95.

20. Liu, J., Zhu, Y., Luo, G.Z., Wang, X., Yue, Y., Wang, X., Zong, X., Chen, K., Yin, H., Fu, Y., et al. (2016b). Abundant DNA 6mA methylation during early embryogenesis of zebrafish and pig. Nat Commun 7, 13052.

21. Luo, G.Z., Blanco, M.A., Greer, E.L., He, C., and Shi, Y. (2015). DNA N(6)-methyladenine: a new epigenetic mark in eukaryotes? Nat Rev Mol Cell Biol 16, 705–710.

22. Ma, M., Zhu, H., Zhang, C., Sun, X., Gao, X., and Chen, G. (2015). “Liquid biopsy”-ctDNA detection with great potential and challenges. Ann Transl Med 3, 235.

23. Mondo, S.J., Dannebaum, R.O., Kuo, R.C., Louie, K.B., Bewick, A.J., LaButti, K., Haridas, S., Kuo, A., Salamov, A., Ahrendt, S.R., et al. (2017). Widespread adenine N6-methylation of active genes in fungi. Nat Genet 49, 964–968.

24. Ping, X.L., Sun, B.F., Wang, L., Xiao, W., Yang, X., Wang, W.J., Adhikari, S., Shi, Y., Lv, Y., Chen, Y.S., et al. (2014). Mammalian WTAP is a regulatory subunit of the RNA N6-methyladenosine methyltransferase. Cell Res 24, 177–189.

25. Ratel, D., Ravanat, J.L., Berger, F., and Wion, D. (2006). N6-methyladenine: the other methylated base of DNA. Bioessays 28, 309–315.

26. Shi, L., Guo, Y., Dong, C., Huddleston, J., Yang, H., Han, X., Fu, A., Li, Q., Li, N., Gong, S., et al. (2016). Long-read sequencing and de novo assembly of a Chinese genome. Nat Commun 7, 12065.

27. Smith, Z.D., and Meissner, A. (2013). DNA methylation: roles in mammalian development. Nat Rev Genet 14, 204–220.

28. Timinskas, A., Butkus, V., and Janulaitis, A. (1995). Sequence motifs characteristic for DNA [cytosine-N4] and DNA [adenine-N6] methyltransferases. Classification of all DNA methyltransferases. Gene 157, 3–11.

29. Touzain, F., Petit, M.A., Schbath, S., and El Karoui, M. (2011). DNA motifs that sculpt the bacterial chromosome. Nat Rev Microbiol 9, 15–26.

30. Vasu, K., and Nagaraja, V. (2013). Diverse functions of restriction-modification systems in addition to cellular defense. Microbiol Mol Biol Rev 77, 53–72.

31. von Meyenn, F., Iurlaro, M., Habibi, E., Liu, N.Q., Salehzadeh-Yazdi, A., Santos, F., Petrini, E., Milagre, I., Yu, M., Xie, Z., et al. (2016). Impairment of DNA Methylation Maintenance Is the Main Cause of Global Demethylation in Naive Embryonic Stem Cells. Mol Cell 62, 848–861.

32. Wacker, D., Stevens, R.C., and Roth, B.L. (2017). How Ligands Illuminate GPCR Molecular Pharmacology. Cell 170, 414–427.

33. Wang, J., Vasaikar, S., Shi, Z., Greer, M., and Zhang, B. (2017). WebGestalt 2017: a more comprehensive, powerful, flexible and interactive gene set enrichment analysis toolkit. Nucleic Acids Res.

34. Wang, K., Li, M., and Hakonarson, H. (2010). ANNOVAR: functional annotation of genetic variants from high-throughput sequencing data. Nucleic Acids Res 38, e164.

35. Westbye, M.P., Feyzi, E., Aas, P.A., Vagbo, C.B., Talstad, V.A., Kavli, B., Hagen, L., Sundheim, O., Akbari, M., Liabakk, N.B., et al. (2008). Human AlkB homolog 1 is a mitochondrial protein that demethylates 3-methylcytosine in DNA and RNA. J Biol Chem 283, 25046–25056.

36. Wion, D., and Casadesus, J. (2006). N6-methyl-adenine: an epigenetic signal for DNA-protein interactions. Nat Rev Microbiol 4, 183–192.

37. Wojtas, M.N., Pandey, R.R., Mendel, M., Homolka, D., Sachidanandam, R., and Pillai, R.S. (2017). Regulation of m6A Transcripts by the 3’-->5’ RNA Helicase YTHDC2 Is Essential for a Successful Meiotic Program in the Mammalian Germline. Mol Cell 68, 374–387 e312.

38. Wu, T.P., Wang, T., Seetin, M.G., Lai, Y., Zhu, S., Lin, K., Liu, Y., Byrum, S.D., Mackintosh, S.G., Zhong, M., et al. (2016). DNA methylation on N(6)-adenine in mammalian embryonic stem cells. Nature 532, 329–333.

39. Xiang, Y., Laurent, B., Hsu, C.H., Nachtergaele, S., Lu, Z., Sheng, W., Xu, C., Chen, H., Ouyang, J., Wang, S., et al. (2017). RNA m6A methylation regulates the ultraviolet-induced DNA damage response. Nature 543, 573–576.

40. Xu, S.H., Huang, J.Z., Xu, M.L., Yu, G., Yin, X.F., Chen, D., and Yan, G.R. (2015). ACK1 promotes gastric cancer epithelial-mesenchymal transition and metastasis through AKT-POU2F1-ECD signalling. J Pathol 236, 175–185.

41. Ye, P., Luan, Y., Chen, K., Liu, Y., Xiao, C., and Xie, Z. (2017). MethSMRT: an integrative database for DNA N6-methyladenine and N4-methylcytosine generated by single-molecular real-time sequencing. Nucleic Acids Res 45, D85–D89.

42. Zhang, G., Huang, H., Liu, D., Cheng, Y., Liu, X., Zhang, W., Yin, R., Zhang, D., Zhang, P., Liu, J., et al. (2015). N6-methyladenine DNA modification in Drosophila. Cell 161, 893–906.

43. Zhang, Y., Liu, T., Meyer, C.A., Eeckhoute, J., Johnson, D.S., Bernstein, B.E., Nusbaum, C., Myers, R.M., Brown, M., Li, W., et al. (2008). Model-based analysis of ChIP-Seq (MACS). Genome Biol 9, R137.

44. Zheng, G., Dahl, J.A., Niu, Y., Fedorcsak, P., Huang, C.M., Li, C.J., Vagbo, C.B., Shi, Y., Wang, W.L., Song, S.H., et al. (2013). ALKBH5 is a mammalian RNA demethylase that impacts RNA metabolism and mouse fertility. Mol Cell 49, 18–29.

