## Supplementary Materials for "N^6^-Methyladenine DNA Modification in Human Genome"

**Supplementary information**

### Supplementary Figures

#### Figure S1

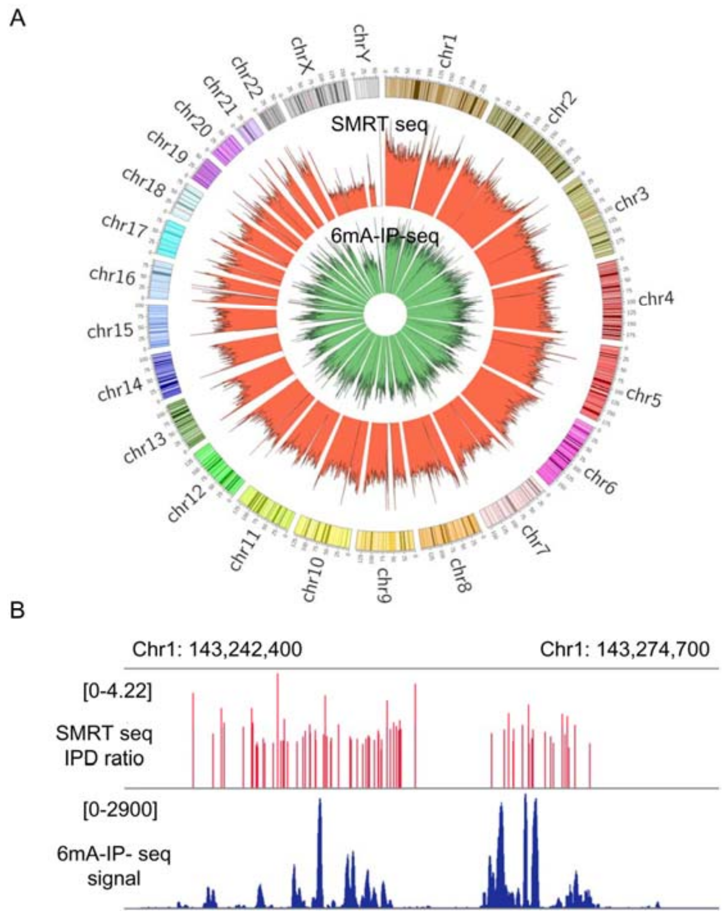


**Figure S1. Related to Figure 1 and 2.** The 6mA sites-occupied regions were highly overlapped between SMRT-sequencing and 6mA-IP-seq in human blood samples. (A) The 6mA-contained DNA fragments identified by 6mA-IP-seq were highly overlapped with the 6mA sites-occupied DNA regions identified by SMRT-sequencing across all chromosomes. (B) Representative overlapping regions of 6mA sites-occupancy between SMRT-sequencing and 6mA-IP-seq.

**Figure S2**

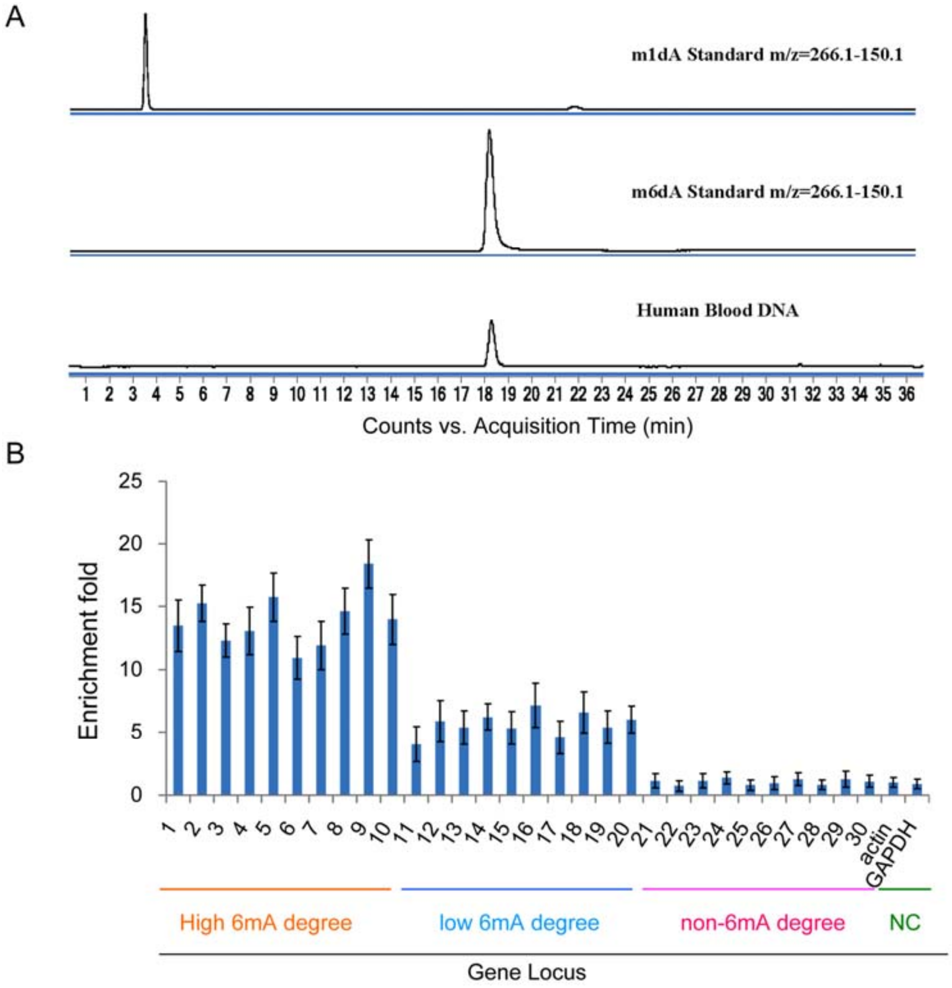


**Figure S2**. **Related to Figure 1.** Validation of 6mA modification in human genomic DNA by LC-MS/MS. (A) LC-MS/MS chromatograms showing 1mA and 6mA levels in the standard oligos and genomic DNA from blood sample. (B) Validation of the 10 gene loci with high 6mA modification, 10 gene loci with low 6mA and 10 gene loci without 6mA from SMRT sequencing by 6mA-IP-qPCR assay (n=3). *Actin* and *GAPDH* was used as a negative control. Results are represented as mean ± SEM.

**Figure S3**

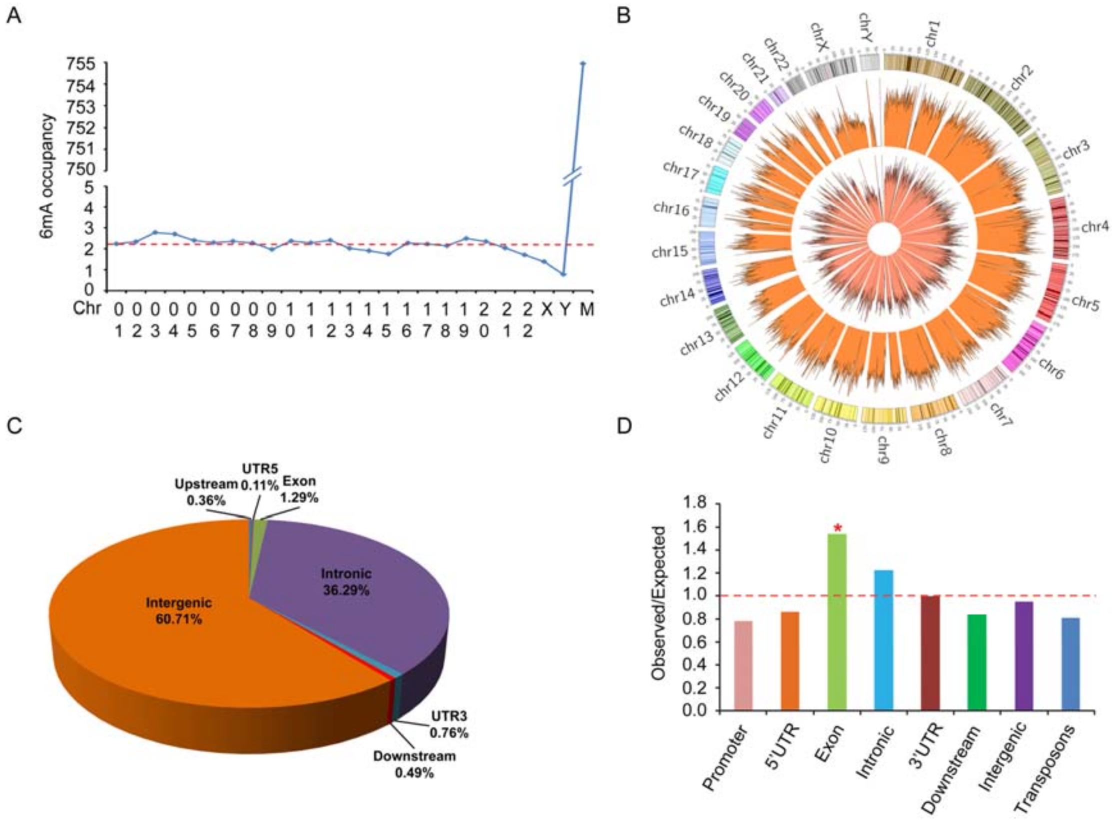


**Figure S3. Related to Figure 1 and 2.** Validation of 6mA modification in human genomic DNA by 6mA-IP-seq, and distribution of the 6mA-containing DNA fragments identified by 6mA-IP-seq in chromosomes and genome. (A) The 6mA-containing DNA fragments across all chromosomes. (B) Circos plots of the 6mA-containing DNA fragments across all human chromosomes in the different 6mA modification level category. (C) The distribution of the 6mA-containing DNA fragments in the functional elements of human genome DNA. (D) Comparison of observed versus expected distributions of the 6mA-containing DNA fragments in each functional elements.

**Figure S4**

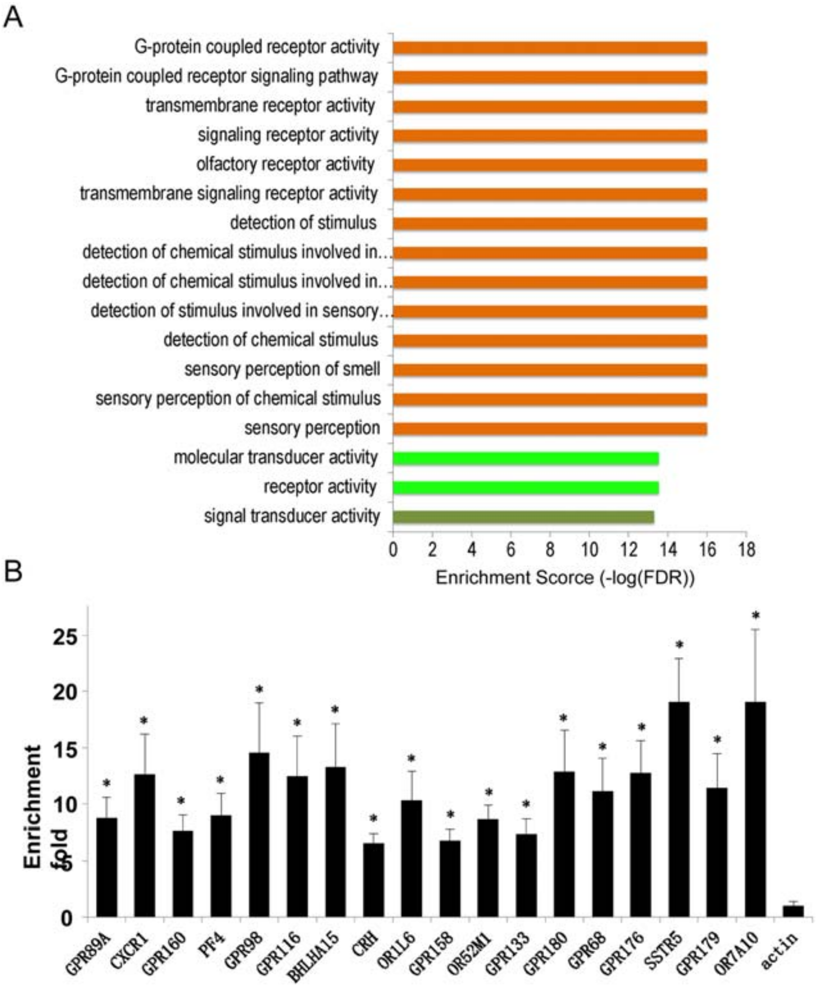


**Figure S4. Related to Figure 1 and 2.** GPCR family gene loci had high 6mA modification (A) GO enrichment category of TOP 200 genes with higher 6mA modification density. (B) Validation of the 18 identified 6mA-methylated GPCR family gene loci from SMRT sequencing by 6mA-IP-qPCR assay (n=3). *Actin* was used as a negative control. Results are represented as mean ± SEM.

**Figure S5**

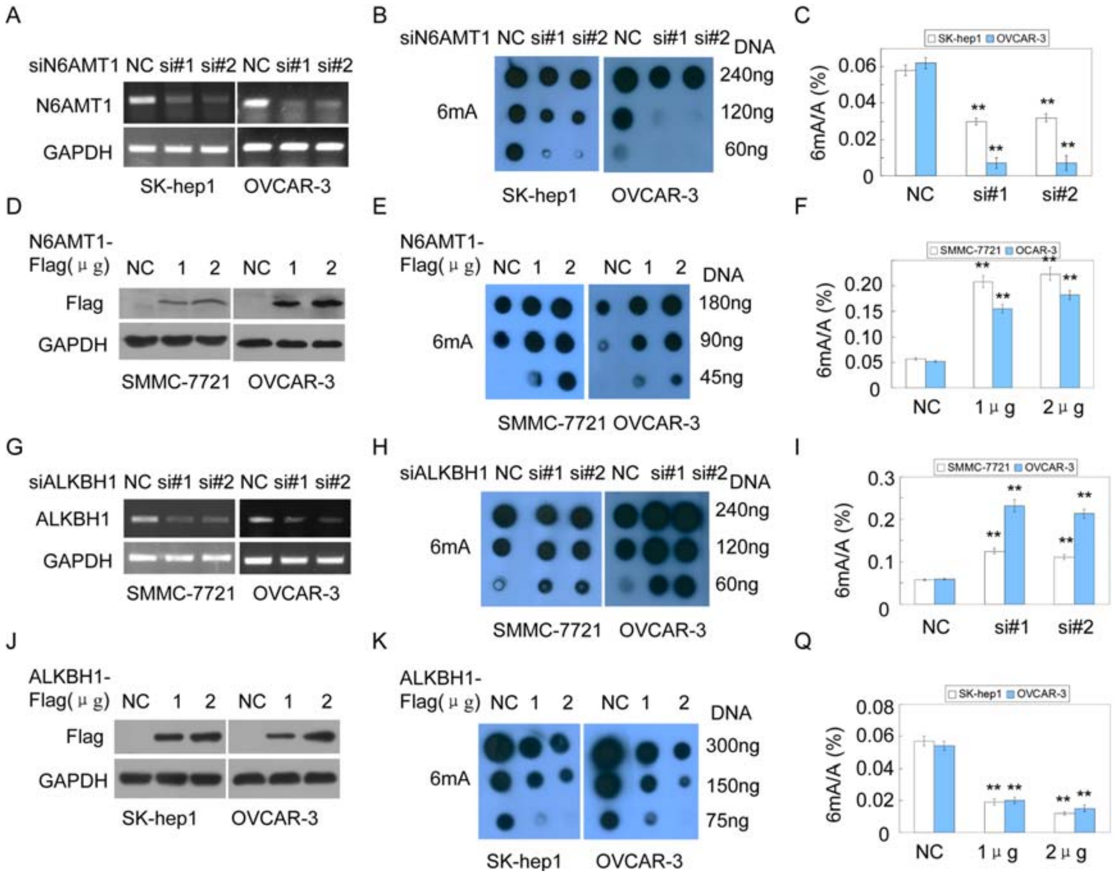


**Figure S5**. **Related to Figure 3 and 4.** N6AMT1 and ALKBH1 are a methyltransferase and demethylase for DNA 6mA methylation and demethylation in human. (A-C) The indicated human cells were transfected with anti-N6AMT1 siRNAs, the N6AMT1 levels detected by RT-PCR (A), the 6mA levels of genomic DNA were subjected to dot blotting assays using a specific anti-6mA antibody (B) and LC-MS/MS assays (n=3) (C). (D-F) The indicated cells were transfected with the indicated amounts of N6AMT1-Flag plasmids, the N6AMT1 levels were determined by western blotting (D), the 6mA levels of genomic DNA were subjected to dot blotting (E) and LC-MS/MS assays (n=3) (F). (G-I) The indicated cells were transfected with anti-ALKBH1 siRNAs, then the ALKBH1 levels were detected by RT-PCR (G), the 6mA levels of genomic DNA were subjected to dot blotting (H) and LC-MS/MS assays (n=3) (I). (J-Q) The indicated cells were transfected with the indicated amounts of ALKBH1-Flag plasmids, then the ALKBH1 levels were determined by western blotting (J), and the 6mA levels of genomic DNA were subjected to dot blotting (K) and LC-MS/MS assays (n=3) (Q). Results are represented as mean ± SEM.

**Figure S6**

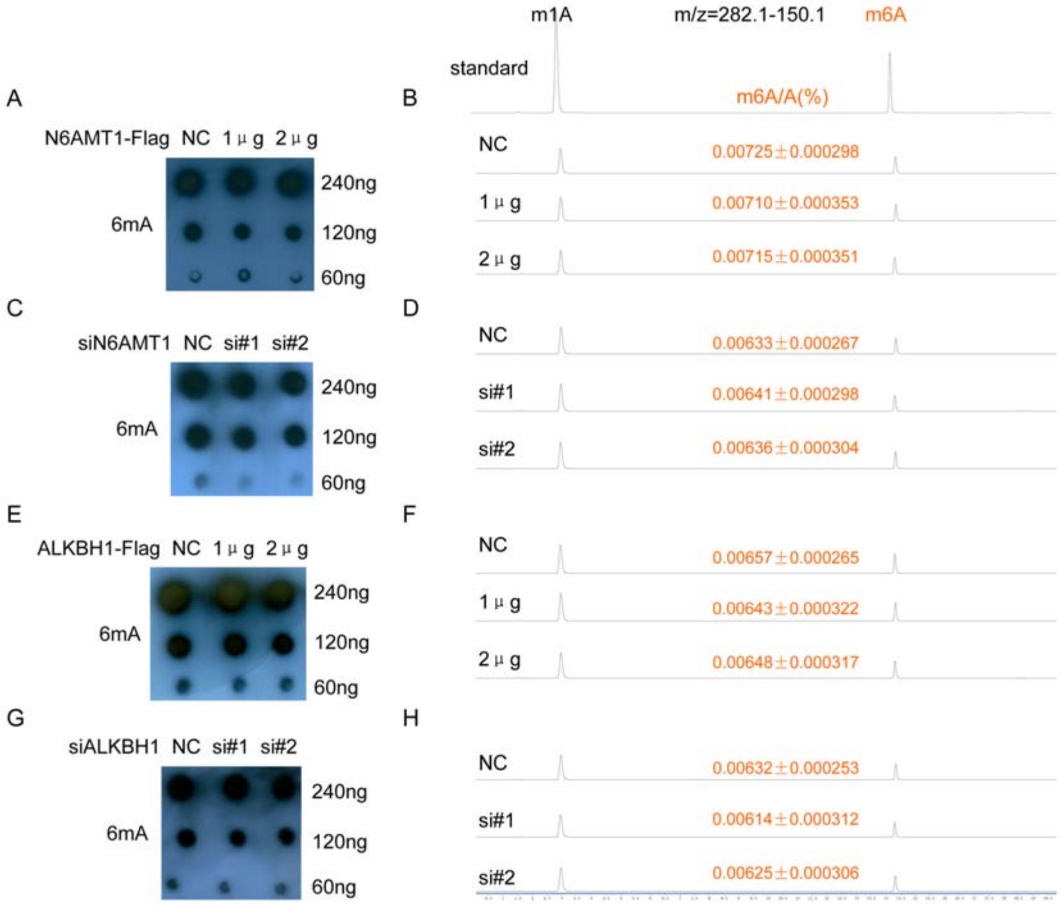


**Figure S6**. **Related to Figure 3 and 4.** N6AMT1 and ALKBH1 did not change the RNA 6mA levels. (A, B) BEL-7402 cells were transfected with the indicated amounts of N6AMT1-Flag plasmids, the RNA 6mA levels were subjected to dot blotting assays using a specific anti-6mA antibody (A) and LC-MS/MS assays (B). (C, D) MHCC-LM3 cells were transfected with anti-N6AMT1 siRNAs, the RNA 6mA levels were subjected to dot blotting (C) and LC-MS/MS assays (D). (E, F) MHCC-LM3 cells were transfected with the indicated amounts of ALKBH1-Flag plasmids, the RNA 6mA levels were subjected to dot blotting (E) and LC-MS/MS (F) assays. (G, H) BEL-7402 cells were transfected with anti-ALKBH1 siRNAs, the RNA 6mA levels were subjected to dot blotting (G) and LC-MS/MS assay (H). Results are represented as mean ± SEM.

### Figure S7

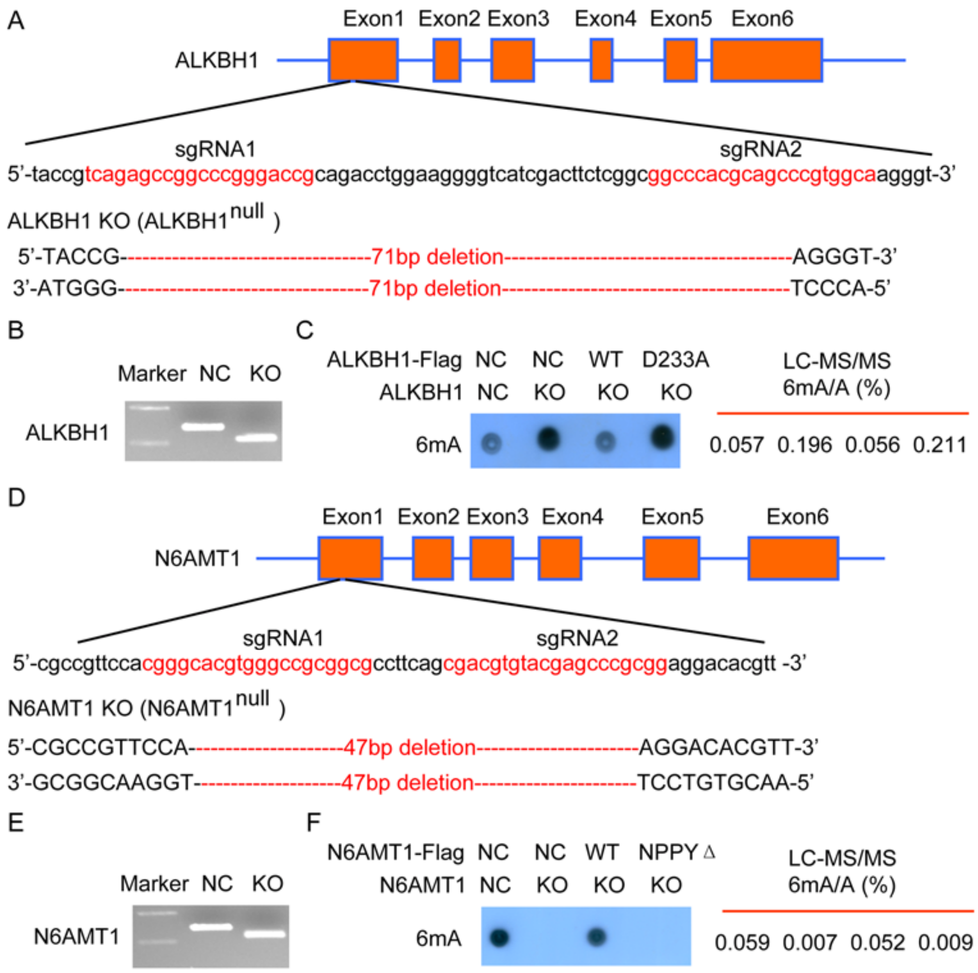


**Figure S7. Related to Figure 3 and 4.** The increase and reduction of genomics DNA 6mA in *ALKBH1* and *N6AMT1* Knockout (KO) HEK293T cells could be efficiently rescued by ectopic expression of wild-type but not mutant *ALKBH1* and *N6AMT1*, respectively. (A, B) Schematic of the *ALKBH1* gene knockout strategy (the Cas9/sgRNAs target sites indicated in red) (A) and confirmed by PCR (B). (C) The indicated ALKBH1 mutant plasmids were transfected into *ALKBH1* KO HEK293T cells, the genomics 6mA levels were determined by dot blotting (left panel) and LC-MS/MS (right panel) assays. (D, E) Schematic of the *N6AMT1* gene knockout strategy (the Cas9/sgRNAs target sites indicated in red) (D) and confirmed by PCR (E). (F) The indicated *N6AMT1* mutant plasmids were transfected into *N6AMT1* KO HEK293T cells, the genomics 6mA levels were determined by dot blotting (left panel) and LC-MS/MS (right panel) assays.

**Figure S8**

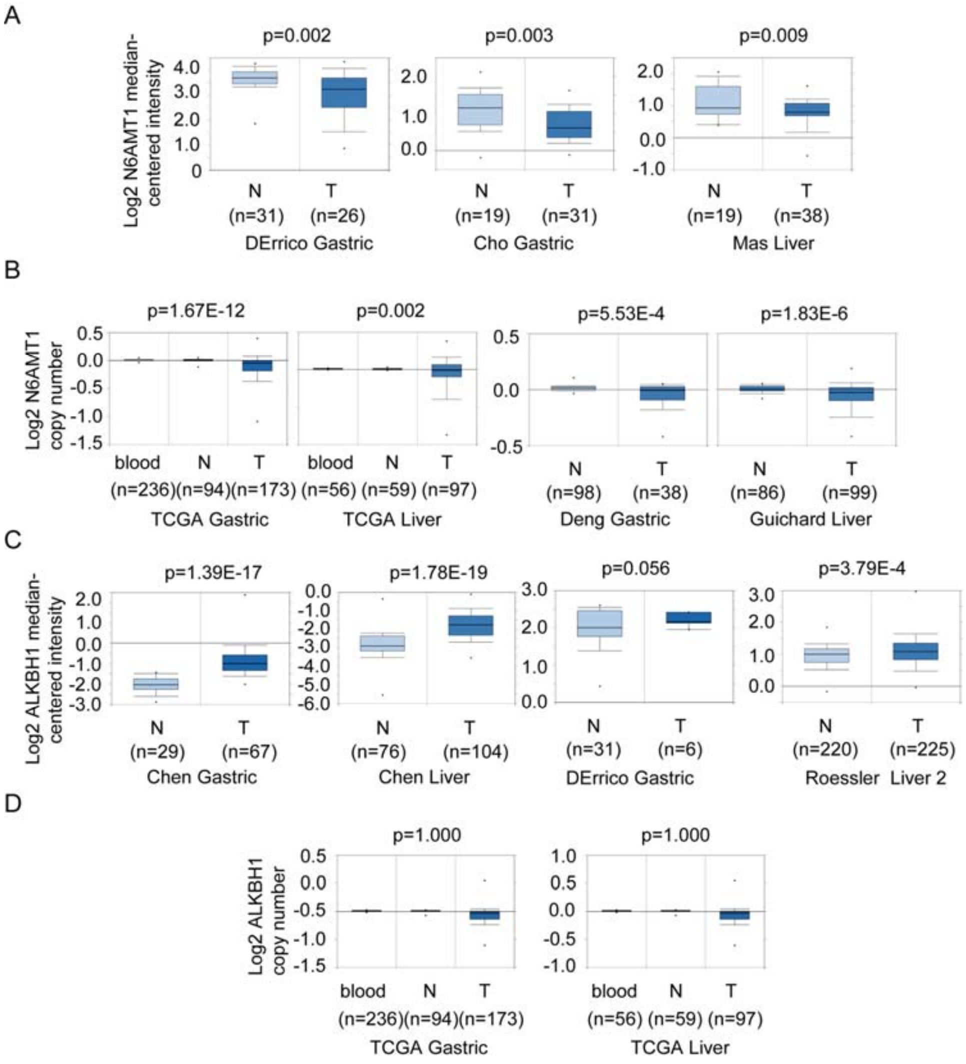


**Figure S8. Related to Figure 5.** N6AMT1 mRNA level and gene copy number were down-regulated, while ALKBH1 mRNA level was up-regulated and its gene copy number was changed in cancer tissues compared to their normal tissues. (A, B) N6AMT1 mRNA level and gene copy number between normal tissues (N) and cancer tissues (GC) was analyzed in the Oncomine gastric and liver cancer database. (C, D) ALKBH1 mRNA level and gene copy number between normal tissues (N) and cancer tissues (GC) was analyzed in the Oncomine gastric and liver cancer database.

**Figure S9**

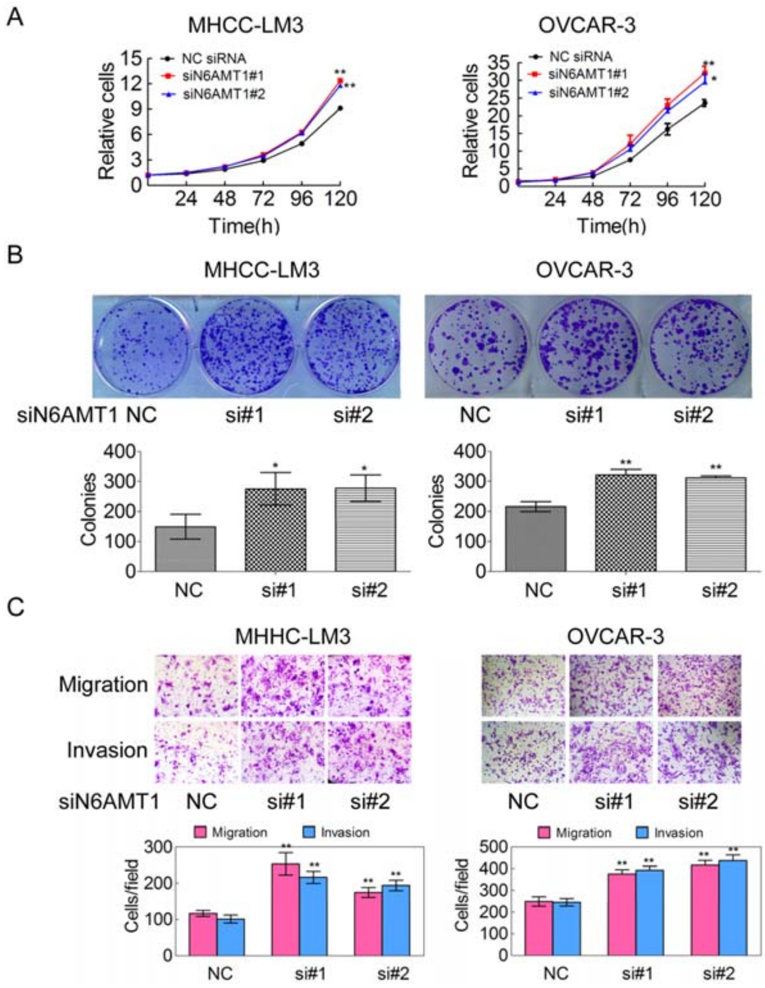


**Figure S9**. **Related to Figure 6.** Silencing of N6AMT1, accompanying reduction of 6mA modification level, promotes tumorigenesis in cancer cells. (A-C) The indicated cancer cells were transfected with anti-N6AMT1 siRNAs, the cell growth (A), colony formation (upper panel in B), migration and invasion (upper panel in C) abilities were determined. The colony number (low panel in B) and migrated and invasive cancer cell number (low panel in C) were counted (n=3). The data are represented as the means ± SEM.

**Figure S10**

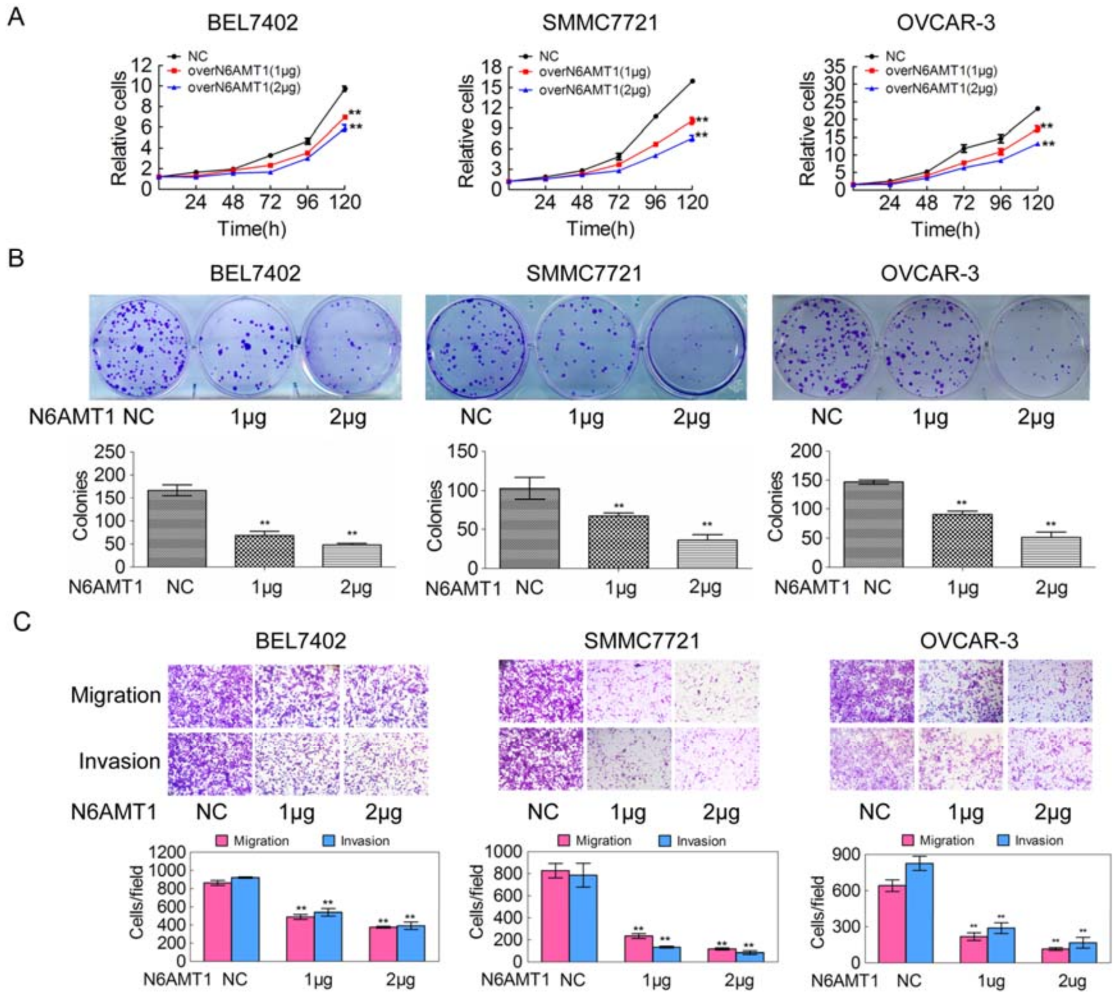


**Figure S10**. **Related to Figure 6.** N6AMT1 over-expression, accompanying increase of 6mA modification level, inhibits tumorigenesis in a dose-dependent manner in cancer cells. The indicated cancer cells were transfected with the indicated amounts of N6AMT1 plasmids, the cell growth (A), colony formation (B), migration and invasion (C) abilities were determined (n=3). The data are represented as the means ± SEM.

**Figure S11**

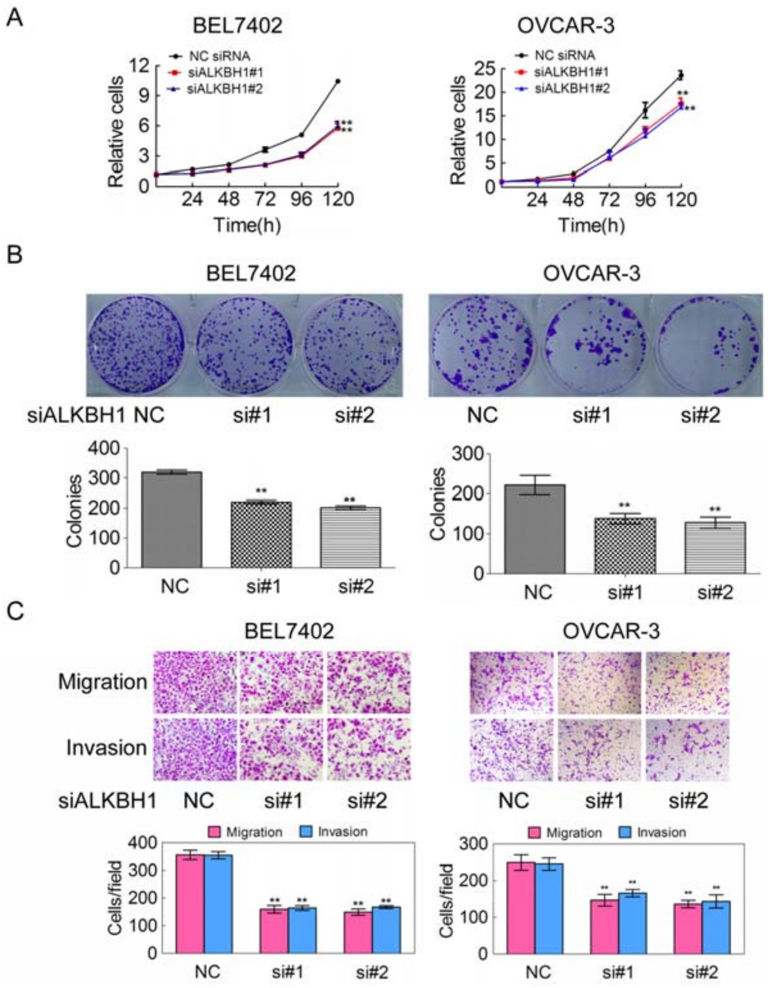


**Figure S11. Related to Figure 7.** Silencing of ALKBH1, accompanying increase of 6mA modification level, inhibits tumorigenesis in cancer cells. (A-C) The indicated cancer cells were transfected with anti-ALKBH1 siRNAs, the cell growth (A), colony formation (B), migration and invasion (C) abilities were determined (n=3). The data are represented as the means ± SEM.

**Figure S12**

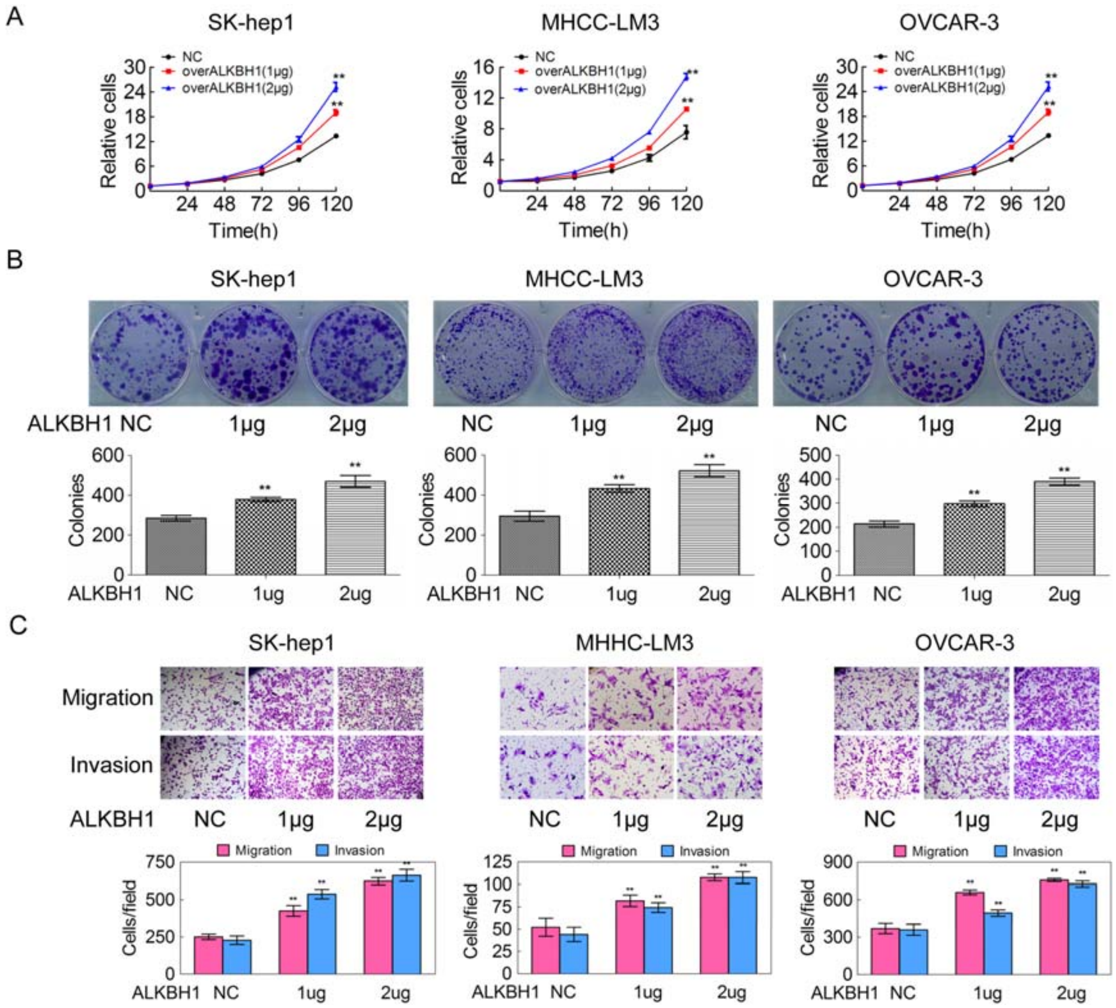


**Figure S12. Related to Figure 7.** ALKBH1 over-expression, accompanying decrease of 6mA modification level, stimulates tumorigenesis in a dose-dependent manner in cancer cells. The indicated cancer cells were transfected with the indicated amounts of ALKBH1 plasmids, the cell growth (A), colony formation (B), migration and invasion (C) abilities were determined (n=3). The data are represented as the means ± SEM.

### Figure S13

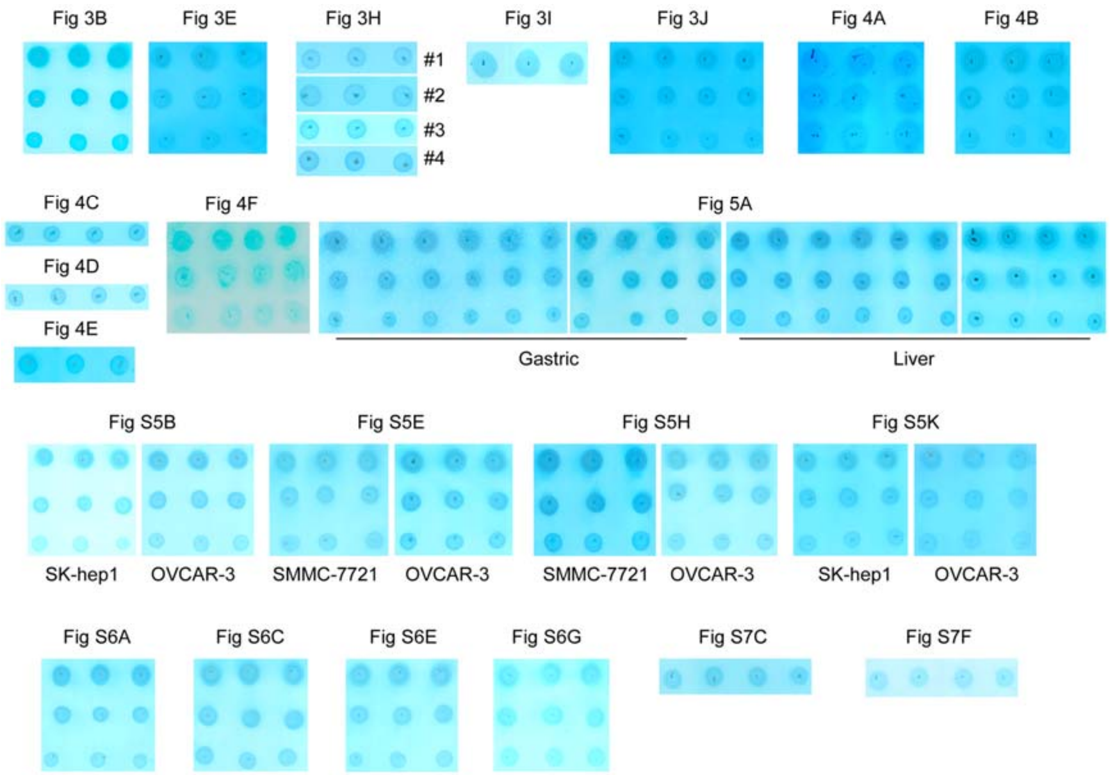


**Figure S13. Related to Figure 3, 4 and 5, and Supplementary Figure S5, S6 and S7.** Loading controls for all 6mA dot blots.

Supplementary Table S6. **Related to Figure 5.** Correlations between DNA 6mA level and clinico-pathological features in 87 liver cancer cases.

| Clinical character | Category | All cases | DNA 6mA# | | $x^2$ | *p* value* |
|---|---|---|---|---|---|---|
| | | | High | Low | | |
| Age (years) | ≤ 60 | 73 | 42 | 31 | 0.001 | 0.978 |
| | > 60 | 14 | 8 | 6 | | |
| Gender | Female | 16 | 10 | 6 | 0.203 | 0.652 |
| | Male | 71 | 40 | 31 | | |
| Tumor size (cm) | ≤3 | 24 | 20 | 4 | 9.069 | 0.003 |
| | >3 | 63 | 30 | 33 | | |
| Cirrhosis | No | 10 | 7 | 3 | 0.726 | 0.394 |
| | Yes | 77 | 43 | 34 | | |
| Histological grade | I-II | 58 | 39 | 19 | 6.795 | 0.009 |
| | III-IV | 29 | 11 | 18 | | |
| AFP (μg/L) | <400 | 32 | 25 | 7 | 8.834 | 0.003 |
| | ≥400 | 55 | 25 | 30 | | |
| Tumor recurrence | No | 51 | 36 | 15 | 8.676 | 0.003 |
| | Yes | 36 | 14 | 22 | | |
| TNM stage | I-II | 59 | 39 | 20 | 5.587 | 0.018 |
| | III-IV | 28 | 11 | 17 | | |
| Tumor number (n) | ≤3 | 83 | 47 | 36 | 0.270 | 0.468 |
| | >3 | 4 | 3 | 1 | | |
| Cirrhosis Nodule size (mm) | ≤3 | 49 | 30 | 19 | 0.647 | 0.421 |
| | >3 | 38 | 20 | 18 | | |

#Nuclear DNA 6mA high: score 3-7; DNA 6mA low: score 0;

**p* value of Fisher's Exact Test;

TNM stage: American Joint Committee on Cancer classification (AJCC).

**Supplementary Methods**

**Antibodies and reagents**

Rabbit anti-N6-methyladenosine antibody was from Millipore (for Dot blotting and IHC, Cat# ABE572). Rabbit anti-N6-methyladenosine antibody was purchased from Synaptic Systems (for 6mA-IP, Cat# 202-003). Mouse anti-DDDDK (Flag)-tag was from MBL (Cat# M185-3L). Mouse anti-β-actin was from Bioworld (Cat# BS6007M).

**Cell culture**

The human hepatocellular carcinoma cell lines BEL-7402, SMMC-7721, MHCC-LM3, SK-hep1 were obtained from the Cell Bank of Shanghai Institutes of Biological Sciences, Chinese Academy of Sciences. The human ovarian cancer cell line OVCAR-3 and kidney cancer cell line HEK293T were obtained from ATCC (USA). All cell lines were tested with Mycoplasma PCR Detection Kit (Sigma-Aldrich). BEL-7402, SMMC-7721 were cultured in RPMI-1640 medium supplemented with 10% fetal bovine serum (FBS), 100 units/ml penicillin and 100 units/ml streptomycin at 37 °C in a 5% $CO_2$ atmosphere, while HCC-LM3, SK-hep1, OVCAR-3 and HEK293T cells were grown in DMEM medium.

**Genome DNA extraction**

Total DNA was extracted from cultured cells and cancer tissues using TIANamp Genomic DNA Kit (Tiangen Biotech, China). To exclude the possibly contaminated methylated-RNA, the extracted genome DNA was treated with 0.1 mg/ml RNase A at 37℃ for 1h. Ultraviolet spectrophotometry was used to detect the content and concentration (A260/A280>1.8) of the genome DNA.

**siRNA**

The cells were transfected with the anti-N6AMT1 or ALKBH1 siRNAs with RNAiMAX (Invitrogen) for 48h. The following siRNA sequences were used: siN6AMT1#1, sense: 5'-GAACUGGCAGGA GUGGAAATTT -3', anti-sense: 5'-UUUCCACUCCUGCCAGUUCTT-3'; siN6AMT1#2, sense: 5'-CCUCAAGUUCACCAAGUCUTT-3', anti-sense: 5'-AGACUUGGUGAACUUG AGGTT-3'; siALKBH1#1, sense: 5'-GCAAGCCUAUGGACUCAAATT-3', anti-sense: 5'-UUUGAGUCCAUAGGCUUGCTT-3'; siALKBH1#2, sense: 5'-GGAAUCCACGUAGACAGAUTT -3', anti-sense: 5'-AUCUGUCUACGUGGA UUCCTT -3'; negative control siRNA, sense: 5'-GCACAAGCUGGAGUACAACU ACATT-3', anti-sense: 5'-UGUAGUUGUACUCCAGCUUGUGCTT-3'.

**Plasmid constructs and transfection**

To generate C-terminal Flag fusion protein constructs with the *N6AMT1* ORF and the *ALKBH1* ORF (N6AMT1-Flag and ALKBH1-Flag), the *N6AMT1* and *ALKBH1* ORF sequences were amplified using RT-PCR and cloned into the pcDNA3.1(+) vector (Invitrogen). These constructs were transfected into cells by Lipo2000 (Invitrogen). The following sequences were used: N6AMT1-Flag tagged, forward: CGGGGTACCAAGGACTATGGCAGGGGAGAAC, reverse: CGGAATTCTTACTTGTCATCGTCGTCCTTGTAGTCAGACTTGGTGAA CTTGAGGACTG; ALKBH1-Flag tagged, forward: CGGGGTACCGCGAGATGGG GAAGATGGCAGC, reverse: CGGAATTCTTACTTGTCATCGTCGTCCTTGTAGT

CGCTGTCAGGGTTTATCCTGGC.

**Construct of synonymous mutant**

The anti-N6AMT1 siRNA-targeted sequences CTGGCAGGAGTGGAA in N6AMT1-Flag and N6AMT1-Flag NPPYΔ mutant plasmids were synonymously mutated to TTAGCCGGCGTAGAG. And the synonymous mutant sN6AMT1-Flag and sN6AMT1-Flag NPPYΔ plasmids were generated.

The anti-ALKBH1 siRNA-targeted sequences CAAGCCTATGGACTCAAA in the ALKBH1-Flag and ALKBH1-Flag D233A mutant plasmids were synonymously mutated to CAGGCATACGGCTTGAAG. And the synonymous mutant sALKBH1-Flag and sALKBH1-Flag D233A plasmids were generated.

**Purification of recombinant proteins**

Confirmed N6AMT1-Flag, N6AMT1-Flag NNPYΔ, ALKBH1-Flag and ALKBH1-Flag D233A plasmids were transfected into HEK293T cells by Lipo2000 (Invitrogen) for 48, respectively. Recombinant N6AMT1-Flag, N6AMT1-Flag NNPYΔ, ALKBH1-Flag and ALKBH1-Flag D233A proteins were purified according to the kit instructions (FLAGIPT1, Sigma). Briefly, anti-Flag M2 affinity resin was added to the cell protein extracts and gently shook on a roller shaker overnight at 4 ºC. The resin was centrifuged for 30s at 5,000g. After three washes with 0.5ml of Wash Buffer, the Flag fusion protein was eluted with 3×Flag peptide. Standard chromatography purification to 95% purity was performed.

**Migration and invasion assays**

*In vitro* migration and invasion assays were performed using Transwell chambers as previously described (Huang et al., 2016; Xu et al., 2015). Briefly, cancer cells were transfected with the indicated plasmids or siRNAs for 36h, respectively. $1\times10^5$ cells in serum-free medium were added in the upper transwell chambers for migration assay (8.0 μM pore size, BD) or the upper transwell chambers coated with Matrigel for invasion assay, and medium supplemented with 10% FBS was added to the bottom chamber. Migrated or invasive cancer cells on the undersurface were stained with 5% crystal violet. Images were captured and the cell number was counted under a microscope.

**Colony formation assays**

Colony formation assays were performed as previously described (Huang et al., 2017). In brief, cancer cells were transfected with the indicated plasmids or siRNAs for 12h. 500 cancer cells were then plated in 6-well culture plates and cultured in medium supplemented with 10% FBS for 2-3 weeks. These cells were then stained with crystal violet solution. The numbers of colonies were counted under the microscope. Three biological replicates were preformed.

**Cellular growth assay**

Cellular growth assays were performed as previously described (Xu et al., 2017). Briefly, cancer cells were transfected with the indicated plasmids or siRNAs for 12h. $1\times10^4$ cancer cells were plated in 96-well culture plates and cultured in medium supplemented with 10% FBS. The cell number was counted at 24, 48, 72, 96, 120h.

**Construct of cell lines with stable silencing of N6AMT1 or ALKBH1**

The lentivirus pLV5-GFP-Luc that expresses green fluorescence protein (GFP) and luciferase (Luc) was purchased from Genepharma (Shanghai, China). SK-hep1 cells were transduced with the lentivirus pLV5-GFP-Luc for 3 days. The Luc-labeled SK-hep1 cells (SK-hep1-Luc) were filtered by flow cytometry and established.

The lentivirus pLV3-N6AMT1 shRNA and pLV3-ALKBH1 shRNA that express anti-N6AMT1 and anti-ALKBH1 shRNA were purchased from Genepharma (Shanghai, China). The SK-hep1-Luc cells were further transduced with the lentivirus pLV3-N6AMT1 shRNA or pLV3-ALKBH1 shRNA for 3 days, and the SK-hep1-Luc-N6AMT1 shRNA and SK-hep1-Luc-ALKBH1 shRNA cells were then selected by using 3μg/ml puromycin for two weeks and established. The silencing of N6AMT1 and ALKBH1 was validated by western blotting.

**Quantitative PCR**

Total RNA was extracted from cells and tissues using the TRNzol Universal Reagent (cat# DP424, Tiangen). RNA reverse transcription was performed by using PrimeScript™RT reagent Kit with gDNA Eraser (cat# RR047A, TAKARA). Quantitative PCR was carried out using SYBR Premix Ex Taq (cat# RR820A, TAKARA). The three independent biological replicates were performed. The primers used here are listed in supplementary Table S7.

### References

Huang, J.Z., Chen, M., Chen, Gao, X.C., Zhu, S., Huang, H., Hu, M., Zhu, H., and Yan, G.R. (2017). A Peptide Encoded by a Putative lncRNA HOXB-AS3 Suppresses Colon Cancer Growth. Mol Cell 68, 171-184 e176.

Huang, J.Z., Chen, M., Zeng, M., Xu, S.H., Zou, F.Y., Chen, D., and Yan, G.R. (2016). Down-regulation of TRPS1 stimulates epithelial-mesenchymal transition and metastasis through repression of FOXA1. J Pathol 239, 186-196.

Xu, S.H., Huang, J.Z., Chen, M., Zeng, M., Zou, F.Y., Chen, and Yan, G.R. (2017). Amplification of ACK1 promotes gastric tumorigenesis via ECD-dependent p53 ubiquitination degradation. Oncotarget 8, 12705-12716.

Xu, S.H., Huang, J.Z., Xu, M.L., Yu, G., Yin, X.F., Chen, D., and Yan, G.R. (2015). ACK1 promotes gastric cancer epithelial-mesenchymal transition and metastasis through AKT-POU2F1-ECD signalling. J Pathol 236, 175-185.

**Supplementary Table S7. Related to Figure S1B, S4B, and Supplementary Methods.** The Primers used for 6mA-IP-qPCR and qPCR.

| Gene name/gene loci | Primers |
|---|---|
| 1 | Sense: TTGGATGTGTAGGGCAATGA<br>Anti-sense: GTGACTGGTGGGATGAAGGT |
| 2 | Sense: GACGACACCTCGCTCACC<br>Anti-sense: GAGGGGAGAAAGTGCAGGAG |
| 3 | Sense: ACATTGACATGGGTGGGTTT<br>Anti-sense: GACACCTTCTCCTGCATGGT |
| 4 | Sense: CAGCAGCATCAGACACATGA<br>Anti-sense: GCTTCACCATCACAGGCTTC |
| 5 | Sense: CTTGTCCGTCAGTGCCAGT<br>Anti-sense: GAGCAGAGGAGCCAGAACAG |
| 6 | Sense: GGGCAAGAGCGAGAGACA<br>Anti-sense: CCCTCGCTCGTTTTCTTTCT |
| 7 | Sense: GCAGGATTCAGGCTATCTGG<br>Anti-sense: AAGGCTCTGAAAAGGGCTTC |
| 8 | Sense: AAGGATCACACGGAGACCAG<br>Anti-sense: CTCTTTCTCCCTGGCCTGT |
| 9 | Sense: CTCACGACCCTCGACTGC<br>Anti-sense: CGTGTTCCACGAGAAGGACT |
| 10 | Sense: CAATGAATCCAGGAGCTGGT<br>Anti-sense: CGGTGGTGATATCCCCTTTA |
| 11 | Sense: TTCCCGTTTCCAAAGACATC<br>Anti-sense: CACAGAGTTGAAACCTCCTATGG |
| 12 | Sense: CCTGGCCAATACAGTGAAGC<br>Anti-sense: AGTGATTGTCCTGCCTCAGC |
| 13 | Sense: GGTTTAGTCTTGGGAGGGTGT<br>Anti-sense: CCGATCCCACAGAAATACAAA |
| 14 | Sense: CAATGAATCCAGGAGCTGGT<br>Anti-sense: GGTGGTGATATCCCCTTTATCA |
| 15 | Sense: CGGTGGTGATATCCCCTTTA<br>Anti-sense: TAATGAATCCAGGAGCTGGTT |
| 16 | Sense: GAGTCCGTCTCCTGGGTGAT<br>Anti-sense: GGTGATGGGGCTTTTGTCTA |

| | |
|---|---|
| 17 | Sense: CTGTGGGATTGGTGGTGATA<br>Anti-sense: TTTTGAAAGGATCAACAAAATTGA |
| 18 | Sense: TCCTCCCCTTTTGATCCTCT<br>Anti-sense: TGTACCAAGCAGAGTACCTTTGA |
| 19 | Sense: CTGTGGGATTGGTGGTGATA<br>Anti-sense: TTGAAAGGATCAGCAAAATTG |
| 20 | Sense: TGGCACCTTCCTTTGGATAC<br>Anti-sense: TAGGCTTCAAGGCGCTCTAA |
| 21 | Sense: TGAAACCAGAGCTTCCCTCA<br>Anti-sense: CTGGAATGAATGAGCAGCAA |
| 22 | Sense: CCTGTCCTGATCACAGTGGA<br>Anti-sense: TTTGGGAGGTTAAAGCAGGA |
| 23 | Sense: GTCAGCAACAACACCCACAC<br>Anti-sense: CATCCTTGCTTCCTCTGGAC |
| 24 | Sense: CTTGACTGTTGGTGGTGCAT<br>Anti-sense: CTCCATCATCTCAACCCAAAA |
| 25 | Sense: CAAAACAGCCACCCTTGAAT<br>Anti-sense: TTACCCTAAGGCCCAAATCC |
| 26 | Sense: GCCCAGTGAAACTGATCTGG<br>Anti-sense: GCCCCATTTTTGTTAGTGTCC |
| 27 | Sense: CCTGGGCAACAAGAGTGAA<br>Anti-sense: TGATTCCTTCTTCGGCAAAC |
| 28 | Sense: CAGGAAGCTCCTGAAAATGTC<br>Anti-sense: GCCTGACACACGGTAGATGA |
| 29 | Sense: TGATGAGCACCTTGTTGGAG<br>Anti-sense: TGGGGCATGATGAGTAACTG |
| 30 | Sense: CAGAAGGATCCCATCCGTTA<br>Anti-sense: CCAACAAACGAGGAGCCTAA |
| GPR89A | Sense: GCAATTGTGAATGGGAGTTCA<br>Anti-sense: GTTGTAGATATGCGGCGTGA |
| CXCR1 | Sense: GGATTCACCCTGCGTACACT<br>Anti-sense: TGAGGACGACAGCAAAGATG |
| GPR160 | Sense: TACTCGGGAAGGCTGAGGTA<br>Anti-sense: AGACGGAGTCTCGCTCTGC |
| PF4 | Sense: TGGTTAGAACAACTTCTGTCT<br>Anti-sense: CTCCATGAGTAGCCAGAGGT |
| GPR98 | Sense: AGGAAGTTGGTTACATGCAG<br>Anti-sense: TCTTGAACTCCTGGACTCAG |

| | |
|---|---|
| GPR116 | Sense: GAGAGGGGCTTTGGAAAATC<br>Anti-sense: AATACGGTCAGGCCAAGGTA |
| BHLHA15 | Sense: CGTGACAGCAGCATCCAG<br>Anti-sense: AGCGTCTCGATCTTGGAGAG |
| CRH | Sense: GAAGACAACCTCCAGAGAAAGC<br>Anti-sense: CAAAGTTGGTGGCGTGTTC |
| OR1L6 | Sense: GCTACCTGCTGGCCTCTATG<br>Anti-sense: CAGGGAATGTAGGTGGGAGA |
| GPR158 | Sense: GGGTATACGCCCAAAGGATT<br>Anti-sense: GATGGACATTTTGGGTTGG |
| OR52M1 | Sense: GGAGGCACTGGGACTAACAG<br>Anti-sense: GCTCCTGAAGTGACCCTTTG |
| GPR133 | Sense: GACGAAGGGCAGCGACAGCGT<br>AACTCCTACCCACCTGCAAA |
| GPR180 | Sense: CAGTGAAACACATTTGAGAATTTTG<br>Anti-sense: ACGTTTGTGGAGACGGAATC |
| GPR68 | Sense: TGCTTCGTCAGCGAGACCAC<br>Anti-sense: AACAGCTCGGGCTCCTCACC |
| GPR176 | Sense: ACCCACAAGTATGTATCCTAC<br>Anti-sense: GATTGTGCCACTGCACGCCAG |
| SSTR5 | Sense: GAGGACAGAGCTGGCTGAAG<br>Anti-sense: TGTGTGGCAGACGGTTAGAG |
| GPR179 | Sense: TGAGGGCTGCACCAGCTGCAT<br>Anti-sense: GCATGCAGCAGGCCTGGCAG |
| OR7A10 | Sense: CTTCCTTCTGACCGTGATGG<br>Anti-sense: CAGAACCAGCAGTCCACAGA |
| actin | Sense: GCTGTTCCAGGCTCTGTTCC<br>Anti-sense: ATGCTCACACGCCACAACATG |
| GAPDH<br>(6mA-IP-qPCR) | Sense: CGACCACTTTGTCAAGCTCA<br>Anti-sense: AGGGGTCTACATCCGAACTG |
| N6AMT1 | Sense: CGGGAAGTCATGGACAGGTT<br>Anti-sense: GGAAAGTGCAGTGGTTCCTTG |
| ALKBH1 | Sense: AAACTGCGTTGGGTGACCGTA<br>Anti-sense: CTTCAGCTCGGAAATCCTCAAATC |
| GAPDH<br>(qPCR) | Sense: GAAGGTGAAGGTCGGAGTC<br>Anti-sense: AAGATGGTGATGGGATTTC |
